# Mapping the dynamics of epigenetic adaptation during heterochromatin misregulation

**DOI:** 10.1101/2023.07.10.548368

**Authors:** Ajay Larkin, Colin Kunze, Melissa Seman, Alexander Levashkevich, Justin Curran, Dionysus Morris-Evans, Sophia Lemieux, Ahmad S. Khalil, Kaushik Ragunathan

## Abstract

A classical and well-established mechanism that enables cells to adapt to new and adverse conditions is the acquisition of beneficial genetic mutations. Much less is known about epigenetic mechanisms that allow cells to develop novel and adaptive phenotypes without altering their genetic blueprint. It has been recently proposed that histone modifications, such as heterochromatin-defining H3K9 methylation (H3K9me), normally reserved to maintain genome integrity, can be redistributed across the genome to establish new and potentially adaptive phenotypes. To uncover the dynamics of this process, we developed a precision engineered genetic approach to trigger H3K9me redistribution on-demand in fission yeast. This enabled us to trace genome-scale RNA and chromatin changes over time prior to and during adaptation in long-term continuous cultures. Establishing adaptive H3K9me occurs over remarkably slow time-scales relative to the initiating stress. During this time, we captured dynamic H3K9me redistribution events ultimately leading to cells converging on an optimal adaptive solution. Upon removal of stress, cells relax to new transcriptional and chromatin states rather than revert to their initial (ground) state, establishing a tunable memory for a future adaptive epigenetic response. Collectively, our tools uncover the slow kinetics of epigenetic adaptation that allow cells to search for and heritably encode adaptive solutions, with implications for drug resistance and response to infection.

## INTRODUCTION

Adaptation enables cells to survive new or changing environments by establishing novel phenotypes that enhance cell fitness.^1,2^ These processes are dynamic and constantly reshape how organisms respond to a wide range of physiological contexts that includes how cells in our body respond to infections, cancer cells react to chemotherapeutic agents, and the emerging threat of antibiotic and antifungal resistance amongst microbes. ^3–5^ One major mechanism that cells leverage to acquire new phenotypes is altering their DNA sequence through genetic mutations. ^6^ Although beneficial mutations in populations are rare, cells that acquire such mutations will eventually outcompete those that fail to adapt.^7,8^ However, genetic mutations represent an inflexible commitment to a new environment that cannot be reversed following a return to cellular homeostasis.^9,10^ Furthermore, it is well known that genetic adaptation to one condition is often associated with a fitness loss in other environments and hence such changes may represent sub-optimal and terminal solutions amidst fluctuating environments.^11^

An alternative is epigenetic adaptation, whereby cells acquire new phenotypes without any changes to their genetic blueprint.^12^ While genetic mutations are irreversible, epigenetic changes can buffer against deleterious mutations without compromising the overall fitness of the cell.^11,13^ In principle, this strategy offers a dynamic, reversible, and flexible form of adaptation well-suited to rapidly changing environmental conditions especially when such conditions persist only for a few generations.^14–16^ Moreover, due to the flexibility of this mode of adaptation, epigenetic changes often pose serious clinical challenges during the evolution of chemotherapy resistance in cancer cells or the widespread emergence of antifungal resistance.^17–21,21^ Thus, understanding how cells leverage adaptive epigenetic mechanisms and targeting such pathways can help us achieve improved clinical outcomes.

One known example of epigenetic adaptation is prion switching in yeast. In particular, the [*PSI*^+^] prion is the aggregated, self-propagating form of the yeast translation-termination factor Sup35.^22,23^ Upon switching to the prion form, the [*PSI*^+^] prion sequesters soluble (active) Sup35, thereby uncovering previously cryptic genetic variation by promoting genome-wide translation readthrough. Hence, a latent, aggregation-prone, conformational state, when unleashed, can enable cells to acquire novel and heritable phenotypes that may be beneficial in unanticipated conditions. Can other epigenetic pathways be similarly leveraged in an on-demand fashion to unravel latent, heritable, and adaptive phenotypes?

Several lines of evidence suggest that cells have the capacity to alter their transcriptomes in response to stress through stochastic changes in transcription, alterations in chromatin accessibility, rewiring existing regulatory networks, and orchestrating wholesale changes in histone modification states.^18,24–30^ Moreover, recent work has shown that diverse histone modifications with other canonical functions may have adaptive potential by being dynamically redistributed to new genomic loci under different stress conditions.^19–21^ How cells exploit these heritable, chromatin-based epigenetic programs to discover genes that can be activated or repressed to enhance fitness and survival remains mysterious.

To achieve successful adaptation, the dynamic redistribution of histone modifications must in principle meet three critical requirements. The first requirement involves spatial changes in modification states, either through spreading from existing sites or the formation of new islands at novel locations in the genome. This is critical for cells to be able to sample which genes to silence or activate. The second requirement demands that the resulting histone modification-dependent changes in gene expression benefit cells in their new environment.^31^ This process ensures that any optimal adaptive solution that cells make is stably maintained within the population. Lastly, the new cell state should be heritable across multiple generations so that cells are prepared to more rapidly respond to a future instance of being exposed to the initiating stress.^32–36^ Thus, to faithfully map epigenetic adaptation pathways, it is necessary to reconstruct these highly dynamic processes and be able to connect genome-wide changes at the RNA and chromatin levels with cell fitness prior to and following adaptation.

To reconstruct these dynamics, we developed an experimental system based on the fission yeast, *Schizosaccharomyces pombe*. In *S. pombe*, H3K9 methylation (H3K9me) specifies silent epigenetic states otherwise referred to as heterochromatin.^37^ Although heterochromatin normally resides at regulatory regions of the genome, such as centromeres and telomeres, H3K9me can also be deployed to downregulate novel targets.^38–46^ One example of an acute stress in *S.pombe* that elicits an adaptive epigenetic response is so-called “heterochromatin misregulation”. Deleting two major H3K9me antagonists – the H3K14 histone acetyltransferase Mst2 and the putative H3K9 demethylase Epe1 – leads to the adaptive silencing of the sole H3K9 methyltransferase, Clr4, suppressing aberrant genome wide H3K9 methylation and restoring fitness.^47^ We reasoned that this system would provide an ideal, minimal, and genetically pliable framework to induce heterochromatin misregulation and unveil the sequence of events that occur prior to adaptation.

Using synthetic biology, we developed a precision genetic approach to trigger and release heterochromatin misregulation on-demand.^48,49^ Taking inspiration from laboratory evolution experiments, which have been powerful in defining genetic adaptations in microbial populations grown under selective pressure, we coupled this ability to induce heterochromatin misregulation with advanced continuous culture methods that allow us to quantify cell fitness in real-time and identify causal genome-wide transcriptional and chromatin-state changes.^50^ Our inducible experimental system is a significant departure from previous studies that focused primarily on beginning and end-state measurements.^21^ By quantifying cell-fitness in yeast populations, we could precisely trace the time evolution of the adaptive silencing program under multiple cycles of heterochromatin stress and recovery. Our approach uncovers how cells can redistribute H3K9me, records network-level changes in transcription, and defines how this dynamic interplay unlocks cryptic epigenetic variation to enable cell survival under conditions of acute stress. In summary, our study captures key features of how cells turn an existing regulatory pathway that normally ensures H3K9 methylation is deposited only at constitutive sites into an adaptive mechanism with implications for drug resistance and response to infection.

## RESULTS

### An inducible Epe1 depletion system to trigger heterochromatin misregulation on-demand

Epe1, a putative H3K9 demethylase, and Mst2, an H3K14 acetyltransferase, have additive roles in regulating *S.pombe* heterochromatin. Deleting both Epe1 and Mst2 leads to acute heterochromatin misregulation, which in turn promotes an adaptive epigenetic response.^47^ We first confirmed previously published results by generating *mst2Δ epe1Δ* cells, which successfully adapted by silencing the H3K9 methyltransferase, Clr4. We measured an approximately ∼4 fold decrease in Clr4 mRNA levels and the establishment of adaptive H3K9me2 at the *clr4+* locus (**Figure S1A-C**).

Since genetic deletions can only provide an endpoint phenotypic output, we sought to design a system for triggering heterochromatin misregulation on-demand by inducibly, rapidly, and thoroughly depleting Epe1. In principle, such a system would enable us to observe in real-time how cells respond to the induction of acute heterochromatin misregulation and dynamically trace adaptive pathways. We designed a system to control Epe1 protein production at two levels: 1) we replaced the endogenous *epe1+* promoter with a thiamine-repressible promoter (*nmt81*) such that addition of thiamine represses mRNA transcription and 2) we fused an auxin inducible degron tag to the C-terminus of Epe1 to trigger protein degradation. We refer to this inducibly degradable Epe1 allele as *epe1^deg^*(**Figure 1A**).^48,49^ Adding thiamine to cells grown in liquid media caused a rapid 8-fold reduction in Epe1 mRNA levels which, when combined with napthaleneacetic acid (NAA; a synthetic auxin analog), led to the absence of any detectable Epe1 protein in as little as 30 minutes (**Figure 1B-C**). Thus, Epe1 depletion rapidly leads to negligible protein and transcript levels in less than an hour of exposure to NAA and thiamine.

**Figure 1.**
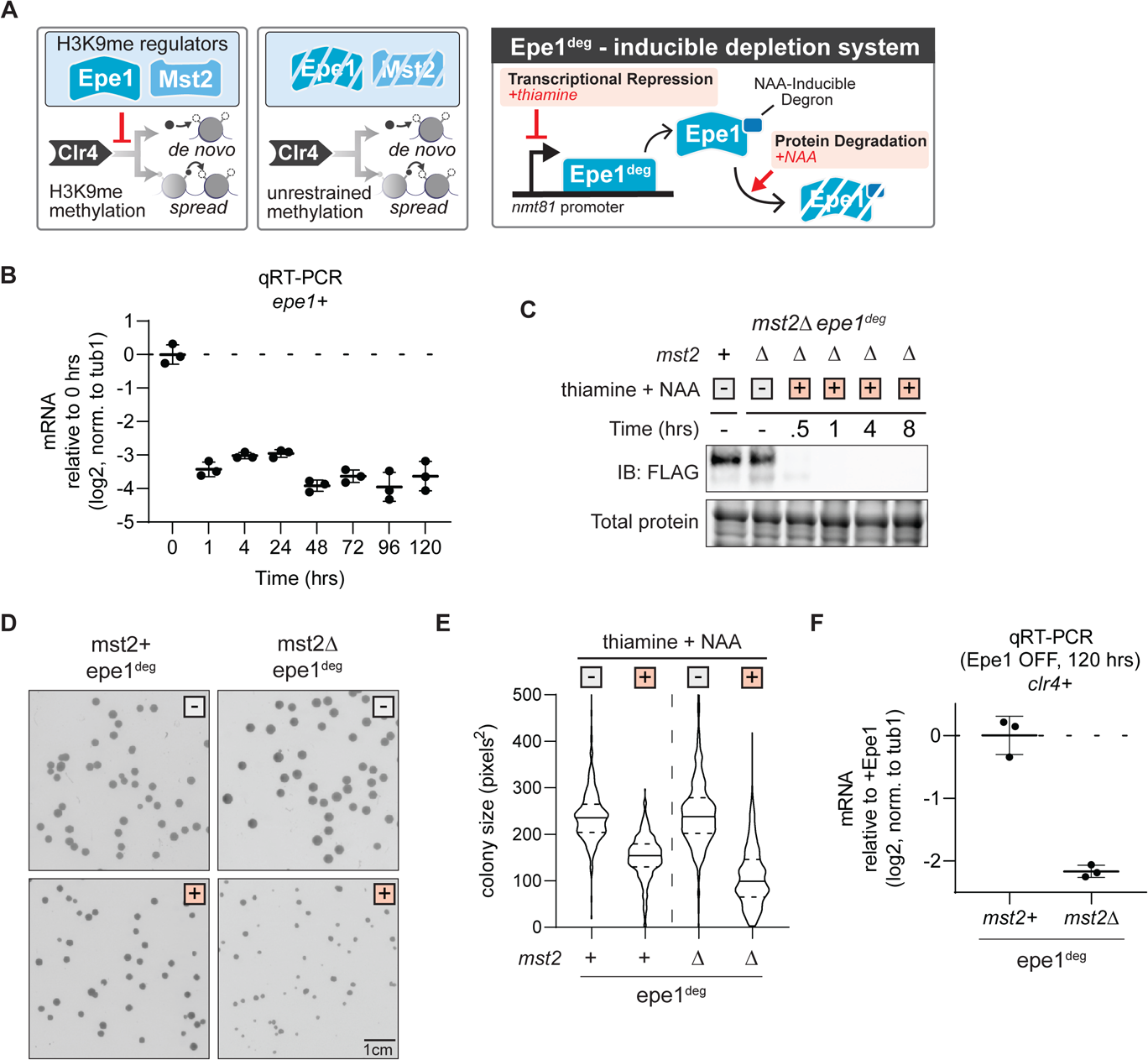
An inducible Epe1 depletion system to trigger heterochromatin misregulation on-demand. (A) Epe1 and Mst2 regulate H3K9me deposition catalyzed by the H3K9 methyltransferase, Clr4 in *S.pombe.* (Left) Epe1 and Mst2 prevent uncontrolled H3K9me spreading. The absence of Mst2 and Epe1 triggers heterochromatin misregulation. (Right) Construction of a precision-engineered genetic approach to toggle Epe1 availability in cells (Epe1^deg^). Epe1 transcription is regulated by a thiamine inducible nmt81 promoter and protein levels are regulated by an auxin-inducible degron tag (AID). Adding auxin and thiamine promotes the on-demand, inducible depletion of Epe1. (B) *epe1+* mRNA expression measured by qRT-PCR as a function of time following treatment with 15µM thiamine and 500µM NAA. Log2 fold-change of mRNA is measured relative to cells without thiamine and NAA (0 hrs). Error bars represent standard deviation, N=3. (C) Western blot for Epe1-3xFLAG-AID in Epe1^deg^ strains. Media type is indicated with either a white box for no treatment, or an orange box for media with 15µM thiamine and 500µM NAA. Total protein levels are shown in the lower panel. (D) Examples of *S.pombe* colonies on solid media after three days of growth. Media type is indicated with either a white box for no treatment, or an orange box for media with 15µM thiamine and 500µM NAA. Image colors are inverted to highlight cell colonies. (E) Colony size distribution measured as pixel area in different genetic backgrounds and growth conditions. Cell size quantified after five days of growth. Media type is indicated with either a white box for no treatment, or an orange box for media with 15µM thiamine and 500µM NAA. Mean and st. dev of distributions in pixels^2^: *mst2+ epe1^deg^* no treatment (240.4 ± 64.2), thiamine and NAA (151.2 ± 46.6); *mst2Δ epe1^deg^* no treatment (246.2 ± 74.6), thiamine and NAA (109.4 ± 64.0) (F) *clr4+* mRNA expression measured by qRT-PCR after five days of treatment with 15µM thiamine and 500µM NAA. Log2 fold-change expression of mRNA is relative to mRNA expression without thiamine and NAA. Error bars represent standard deviation, N=3.

We grew cells overnight and plated equal numbers on non-selective media (white ‘-‘ square) or media that contained NAA and thiamine (orange ‘+’ square) and quantified the mean and standard deviation for colony sizes (**Figure 1D**). We observed generally smaller colonies and substantial colony size heterogeneity when we depleted Epe1 in an *mst2Δ* background, reflecting a fitness loss associated with stress (**Figures 1D-E**). Furthermore, *mst2Δ epe1^deg^* cells exhibited a 4-fold decrease in Clr4 mRNA levels after five days of Epe1 depletion, which recapitulates the adapted state we observed in *mst2Δ epe1Δ* cells (**Figure 1F, S1C**). In contrast, depleting Epe1 in an *mst2+* background produced a less pronounced growth defect and no detectable adaptive Clr4 silencing. Hence, despite the absence of Epe1, these cells exhibited no obvious phenotypic change. Furthermore, there was no decrease in Clr4 mRNA for *mst2+ epe1^deg^* cells after five days of Epe1 depletion, consistent with previous studies.^47^

To test if adaptation was dependent on the order in which the two heterochromatin regulators were depleted, we inverted our genetic background. We developed a strain to deplete Mst2 (*mst2^deg^*) in an *epe1Δ* background: *mst2^deg^ epe1Δ* (**Figure S1D-G**). While there was still some decrease in colony size upon Mst2 depletion, this strain did not produce the same degree of heterogeneity in colony size (**Figure S1E-F**). Additionally, we observed that Clr4 was silenced to a lesser degree compared to *mst2Δ epe1^deg^* cells (**Figure S1G**). Finally, we also developed strains where both Mst2 and Epe1 could be simultaneously depleted in an inducible manner (*mst2^deg^ epe1^deg^*) which would enable us to test if pre-deleting Epe1 or Mst2 produces differences in adaptive phenotypes (**Figure S1H**). The *mst2^deg^ epe1^deg^*exhibited comparable levels of colony size variegation and more robust *clr4+* mRNA suppression compared to *mst2Δ epe1^deg^* (compare **Figure 1C-E** to **Figure S1I-K**).

Nevertheless, even in these strains we noted a residual level of Mst2 protein that remained refractory to depletion after NAA and thiamine addition which we reasoned could potentially have unintended consequences on our adaptation measurements. Collectively, our results establish a system for the inducible, rapid, and complete depletion of Epe1, and demonstrate that the *epe1^deg^* allele recapitulates how *S.pombe* cells adapt in response to acute heterochromatin misregulation.

### Time evolution of adaptive silencing during heterochromatin misregulation

To trace the time evolution of adaptation following Epe1 depletion, we deployed the automated eVOLVER continuous culture platform (**Figure 2A**).^51,52^ eVOLVER enables long-term maintenance of independent *S.pombe* cultures in miniature bioreactors using a continuous turbidostat routine with real-time growth rate quantification.^53–55^ The eVOLVER system also features the ability to schedule media changes, including switching between non-inducer and inducer media. As a result, we can precisely quantify changes in growth resulting from Epe1 depletion, and sample cells as a function of time for molecular measurements to reconstruct the dynamics of Clr4 silencing and concomitant changes in transcription.

**Figure 2.**
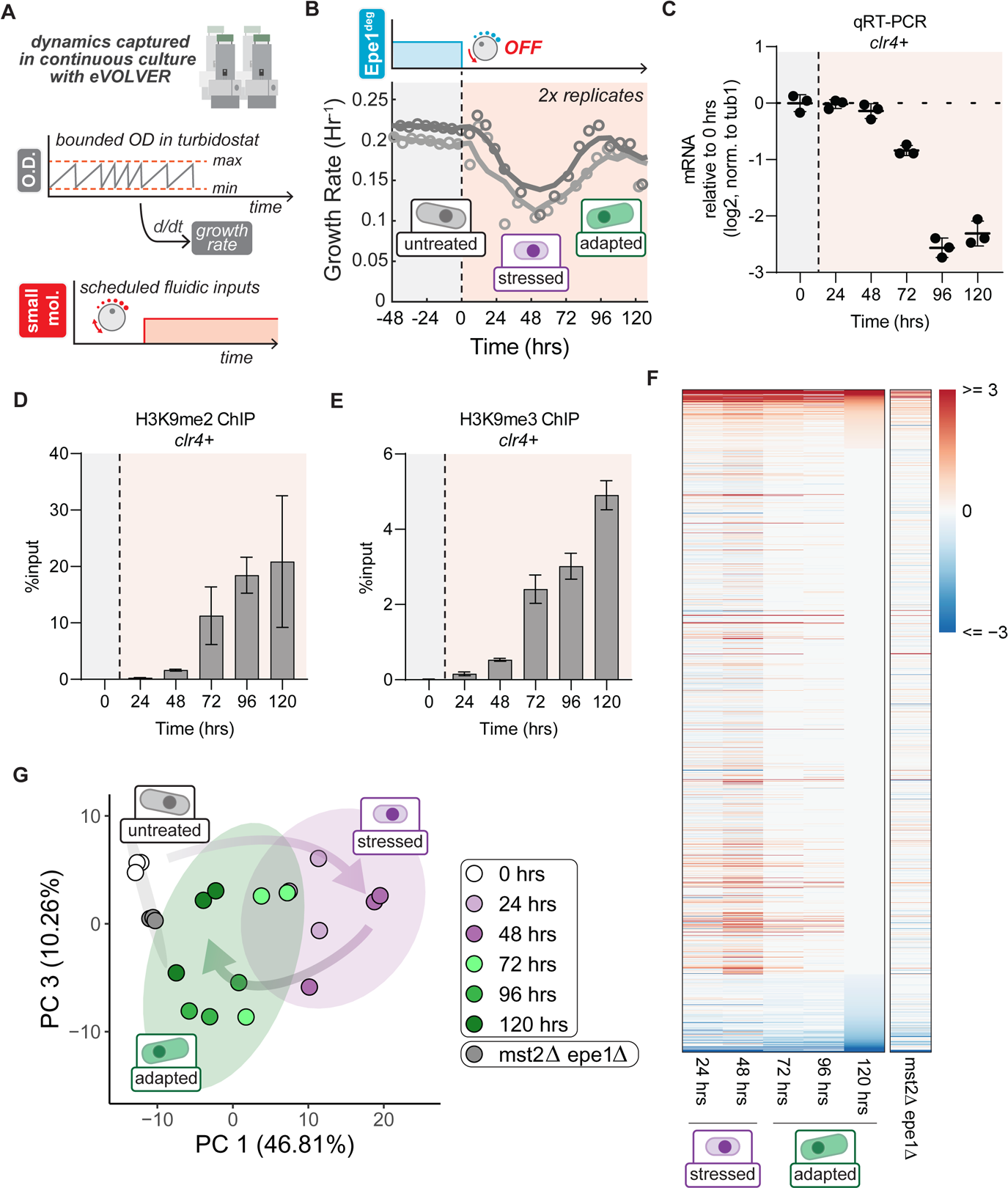
Time evolution of adaptive silencing during heterochromatin misregulation. (A) Model of the eVOLVER continuous culture system used to control growth of *mst2Δ epe1^deg^*cells using a continuous turbidostat routine, with real-time quantification of growth rate and the ability to schedule media changes. (B) Real-time monitoring of growth rates of *mst2Δ epe1^deg^* in eVOLVER. Treatment with 15µM thiamine and 500µM NAA was initiated at t=0hrs. Individual trendlines indicate replicates (N=2). Orange shaded portion represents the time period during which Epe1 has been depleted. (C) *clr4+* mRNA expression measured by qRT-PCR as a function of time following treatment with 15µM thiamine and 500µM NAA. Log2 fold-change of mRNA is measured relative to cells without thiamine and NAA (0 hrs). Orange shaded portion represents the time period during which Epe1 has been depleted. Error bars represent standard deviation, N=3. (D) H3K9me2 ChIP-qPCR measured at the *clr4+* locus as a function of time following treatment with 15µM thiamine and 500µM NAA. The orange shaded portion represents the time period during which Epe1 has been depleted. Error bars represent standard deviation, N=2. (E) H3K9me3 ChIP-qPCR measured at the *clr4+* locus as a function of time following treatment with 15µM thiamine and 500µM NAA. Orange shaded portion represents the time period during which Epe1 has been depleted. Error bars represent standard deviation, N=2. (F) Heatmap of significant differentially expressed genes following treatment with thiamine and auxin relative to untreated *mst2Δ epe1^deg^* cells. Heatmap consists of genes that are differentially expressed at least during one time point. Total number of differentially expressed transcripts N=3896, significance cutoff of AdjPval ≤ 0.01. (G) Time course PCA analysis of the regularized log transform of RNAseq normalized counts denoting different time points after treatment with15µM thiamine and 500µM NAA. Colors denote untreated, stress and adapted cell phases. N=3, ellipse level=0.9.

We grew replicate populations of *mst2Δ epe1^deg^* cells in eVOLVER turbidostats for 48 hours at 32°C before switching to inducer (NAA and thiamine-containing) media to trigger Epe1 depletion (Methods). Upon induction, we observed a substantial reduction in growth rate over the course of a 48-hour period (**Figure 2B**). This was followed by a recovery period in which cells returned to pre-depletion growth rates, indicative of adaptation to heterochromatin misregulation. Based on the eVOLVER time traces, we posited that cells transit through three primary phases upon experiencing heterochromatin misregulation, namely 1) **untreated** (before inducing Epe1 depletion) 2) **stress** (post-induction, characterized by poor growth) and, 3) **adapted** (growth recovery). Replicate eVOLVER populations of *mst2^deg^ epe1^deg^* closely followed the same growth trends as that of the *mst2Δ epe1^deg^* strains (**Figure S2A**). In contrast, we observed little change in the growth rate in *mst2Δ epe1^deg^ clr4Δ* populations upon induction of Epe1 depletion suggesting that the growth rate changes in *mst2Δ epe1^deg^* strains was dependent on H3K9 methylation (**Figure S2B**).

To reconstruct time-resolved changes in Clr4 silencing, we harvested *mst2Δ epe1^deg^* cells grown in 24-hour time intervals for quantification of *clr4+* mRNA and H3K9me2/me3 levels. In the initial stress phase (during the first 48 hours post-induction), we observed no changes in *clr4+* mRNA and very minimal increases to H3K9me2/me3 levels (**Figures 2C-E**). However, during the adapted phase (after 48 hours) we observed a substantial decrease in *clr4+* mRNA expression (**Figure 2C**). The change in *clr4+* mRNA levels coincided with enrichment of H3K9me2/3 at the *clr4+* locus, which had remained largely unmarked until that time (**Figure 2C-E, S2C-D**). Thus, the transition between the stress and adapted phases is closely aligned with a steady enrichment of H3K9me2/me3 and reduction in *clr4+* mRNA levels. These results demonstrate that growth rate and Clr4 silencing dynamics are closely coordinated as cells adapt to acute heterochromatin misregulation stress.

To assess transcriptome-wide changes during the time-course of adaptation, we performed RNA-seq on *mst2Δ epe1^deg^* samples that were collected at 24-hour intervals in triplicate. In the stress phase, we observed acute changes to the transcriptome relative to untreated *mst2Δ epe1^deg^*cells (**Figure 2F**). These gene expression changes during the stress phase gradually vanished by the end of the time course such that, by the time adaptation was completed, the transcriptome of Epe1-depleted cells resembled those of untreated cells. We applied principal component analysis (PCA) to further investigate the transcriptome changes. PCA clearly captured time-dependent transitions between different growth phases following Epe1 depletion (**Figure 2G, S2E**). Notably, gene expression changes in the stress phase (24-48 hours) and those in the late adapted phase (96-120 hours) cluster into non-overlapping statistically significant groups, with the 72-hour time point falling at the intersection between these two groups.

We additionally performed RNA-seq analysis on *mst2Δ epe1Δ* cells to compare with *mst2Δ epe1^deg^* cells. Most strikingly, the transcriptomes of these cells most closely resembled untreated *mst2Δ epe1^deg^* cells (0 hours) (**Figure 2G, S2E**). Importantly, we confirmed that independent *mst2Δ epe1Δ* clones have few differences in their transcriptomes, implying that independent isolates also make the same adaptive choices and their gene expression networks are rewired in a very similar manner (**Figure S2F-G**). *mst2Δ epe1Δ* cells have silenced *clr4+* and have been grown well beyond 120 hours. Thus, their convergence towards the untreated transcriptome implies that there are additional RNA level changes that occur beyond our 120-hour adaptation time-course. For example, we found that, in Epe1-depleted cells at 120 hours, genes associated with iron homeostasis are upregulated while genes associated with ATP synthesis and cellular respiration are downregulated (**Figure S2H-I**), whereas in *mst2Δ epe1Δ* cells these genes returned to expression levels equivalent to untreated *mst2Δ epe1^deg^*cells (**Figure S2J**).^39^ Taken together, our system reveals distinct population-level cell states during the adaptation process.

### Heterochromatin misregulation triggers the targeted expansion of pre-existing H3K9me3 islands

To investigate how heterochromatin misregulation drives changes in the H3K9 methylome over time, we cultured *mst2Δ epe1^deg^* cells as batch cultures over six 24-hour periods encompassing the untreated, stress, and adapted phases. We then performed chromatin immunoprecipitation followed by deep sequencing (ChIP-seq) to map changes in the H3K9 methylome. We observed expansion of specific H3K9me3 domains, primarily at constitutive heterochromatin (pericentromeres, telomeres, and the ribosomal DNA locus) and several heterochromatin islands centered around meiotic genes and ncRNAs (**Figure 3A, Figure S3A, Table S4**).^46,47,56^ We used K-means clustering to separate H3K9me3 peaks into four statistically defined groups. The first group uniquely corresponds to the *clr4+* locus where H3K9me3 is established and maintained throughout the time course of adaptation. The other three groups contain almost all the identified islands, which show a pattern of growth up to 48 hours, then decay by 120 hours while *clr4+* (depicted in group 1) undergoes silencing. We observed very little enrichment for H3K9me2 or H3K9me3 peaks outside of these islands or constitutive heterochromatin. Expansion of H3K9me3 reached a maximum at the end of stress phase (48 hours) followed by a steady decay in the adapted phase at constitutive heterochromatin, non-coding RNAs, and meiotic genes (**Figure 3B-C**). In contrast, H3K9me3 is deposited *de novo* at the *clr4+* locus and accumulates over time (**Figure 3D**). This process is distinct from other H3K9me3 peaks at meiotic genes or ncRNA, which expand during stress and then subsequently retract once cells adapt (**Figure 3E, S3B**). We also validated that the pre-deletion of Mst2 did not drive any pre-adaptation by performing ChIP-seq measurements of H3K9me3 in *mst2^deg^ epe1^deg^* cells, wherein the islands that form during and after adaptation are identical to *mst2Δ epe1^deg^*cells (**Figure S3C-D**).

**Figure 3.**
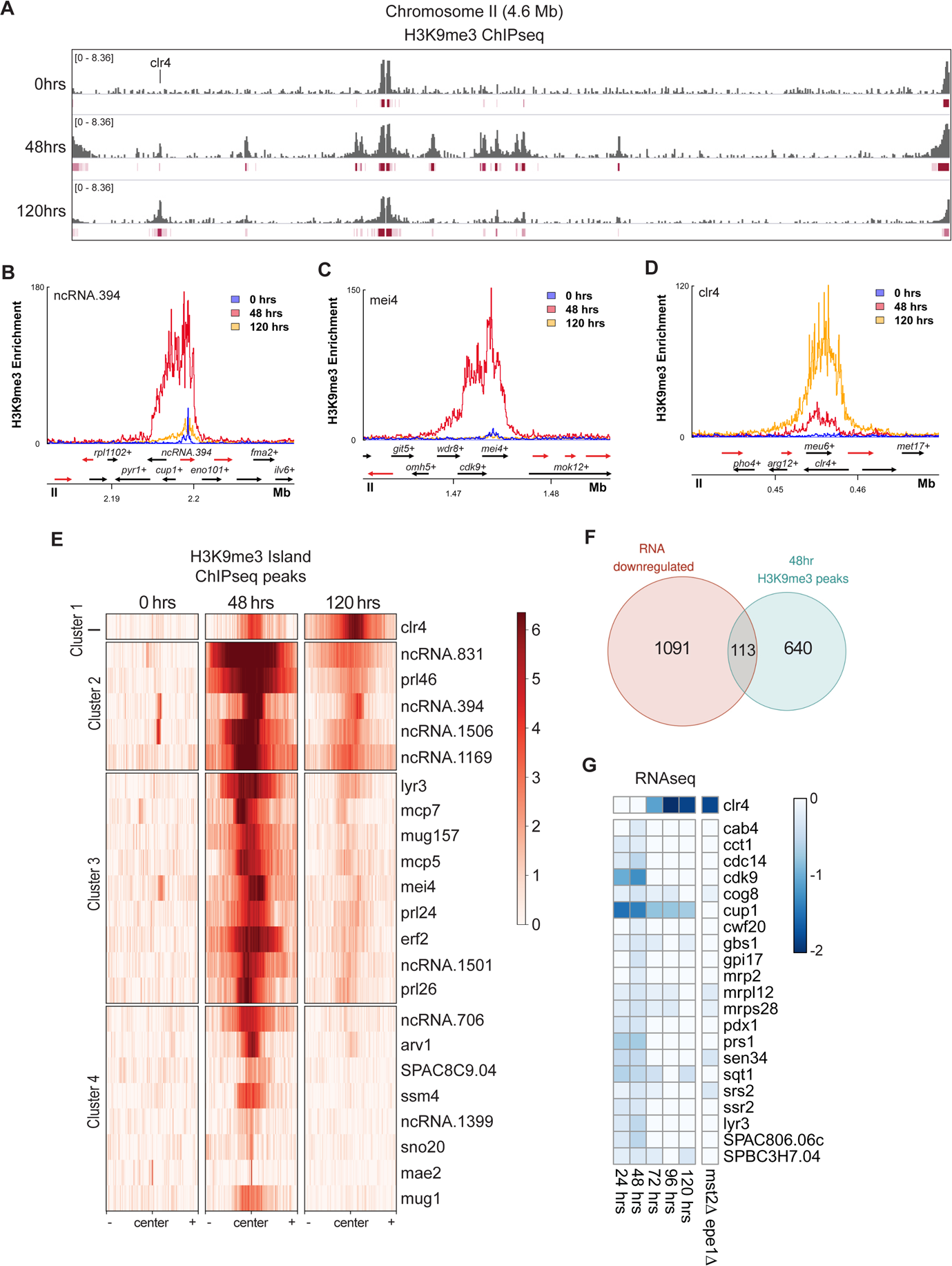
Heterochromatin misregulation triggers the targeted expansion of pre-existing H3K9me3 islands. (A) H3K9me3 ChIP-seq tracks of chromosome II of the *S. pombe* genome. Enrichment is shown in log2 fold change of IP normalized to input. Time of Epe1 depletion is indicated on the left side of each track. Peaks identified are denoted in red below each track. The *clr4+* gene locus is specifically highlighted. One of two ChIP-seq replicates is shown in this figure. (B) H3K9me3 ChIP-seq enrichment centered on *ncRNA.394* for the indicated time points. Genomic tracks below show coding transcripts in black, non-coding transcripts in red. (C) H3K9me3 ChIP-seq enrichment centered on *mei4+* for the indicated time points. Genomic tracks below show coding transcripts in black, non-coding transcripts in red. (D) H3K9me3 ChIP-seq enrichment centered on *clr4+* for the indicated time points. Genomic tracks below show coding transcripts in black, non-coding transcripts in red. (E) K-means clustered heatmap (k=4) of H3K9me3 islands during heterochromatin misregulation at 0hrs, 48hrs, and 120hrs of Epe1 depletion in *mst2Δ epe1^deg^*. Peaks shown in 24kb windows. (F) Venn diagram depicting genes that are downregulated (AdjPval ≤ 0.01) by 48 hours after Epe1 depletion, overlapped with genes marked by H3K9me3 selectively at 48 hours. (G) Heatmap depicting the expression dynamics of essential genes selectively marked by H3K9me3 at 48 hours. Changes in expression are log2 fold change relative to untreated *mst2Δ epe1^deg^* cells. (AdjPval ≤ 0.01)

We cross-referenced our H3K9me3 ChIP-seq and transcriptome time-course data to measure transcriptomic changes caused by aberrant H3K9me spreading during heterochromatin misregulation. We found a total of 753 genes under expanded H3K9me3 peaks during stress phase (48 hours) that were previously not marked by H3K9me3 in the untreated population (**Figure S3E**). Surprisingly, of these 753 genes, a subset of only 113 genes were significantly downregulated (**Figure 3F**). This subset of genes was not functionally enriched for any specific pathways by GO analysis, but notably included a collection of 21 essential genes, including the mitochondrial LYR protein *cup1+.*^21,57^ These essential genes are repressed up until the end of stress phase (48 hours) after which *clr4+* silencing and growth rate recovery coincides with their de-repression (**Figure 3G**). This observation suggests that the downregulation of *cup1+*, and other essential genes proximal to expanding H3K9me3 islands, may correlate with poor cell growth during early heterochromatin misregulation. In contrast, during the adapted phase, there was a dramatic shift in the H3K9me3 methylome. Genes that were marked by novel H3K9me3 and significantly downregulated were proximal to the *clr4+* locus (**Figure S3F-H**). Together, these results indicate that heterochromatin misregulation drives very targeted expansion of existing H3K9 methylation domains, silencing only a small fraction of essential genes. Additionally, development of facultative heterochromatin over the *clr4+* locus occurs *de novo* and represents a rare example of a new ectopic site of H3K9 methylation distinct from the targeted expansions of existing sites of H3K9 methylation.

### Activation of the cellular stress response pathway is required for survival but not adaptive choice

To identify gene pathways relevant to the stress phase of heterochromatin misregulation, we analyzed the set of differentially expressed genes within the stress phase of *mst2Δ epe1^deg^* cells, envisioning that it is most likely to contain the most critical population-level transcriptomic features required for adaptation. Enriched GO terms in this set included genes involved in ribosome biogenesis, translation, caffeine and rapamycin treatment, nitrogen depletion, and the core environmental stress response (CESR) (**Figure 4A, S4A**) ^58,59^. These results were surprising, given our prior assumption that misregulation of the epigenome is a unique form of stress distinct from other types of environmental stresses.

**Figure 4.**
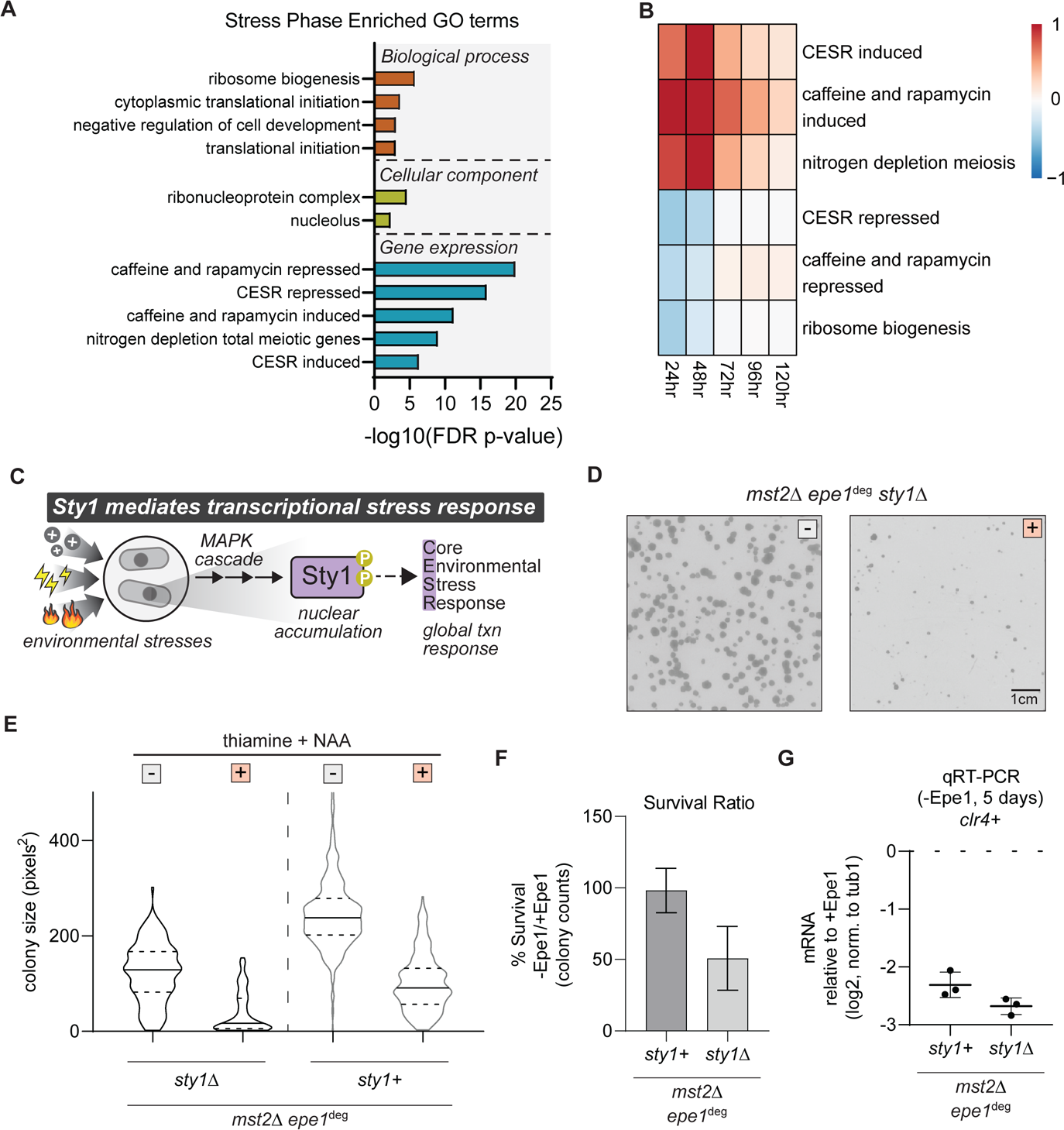
Activation of the cellular stress response pathway is required for survival but not adaptive choice. (A) Selected GO terms for genes differentially expressed (AdjPval ≤ 0.01) within stress phase (48 hours of Epe1 depletion). GO significance cutoff was set as FDR p-value ≤ 0.01. (B) Heatmap showing average fold change for genes differentially expressed (AdjPval ≤ 0.01) in selected GO categories, relative to untreated *mst2Δ epe1^deg^* cells. (C) Environmental stresses trigger a stress-activated MAPK cascade that phosphorylates Sty1, which drives a global transcriptional response that includes the core environmental stress response. (D) Examples of *mst2Δ epe1^deg^ sty1Δ S.pombe* colonies on solid media after three days of growth. Media type is indicated with either a white box for no treatment, or an orange box for media with 15µM thiamine and 500µM NAA. Image colors are inverted to highlight cell colonies. (E) Colony size distribution, in pixel area, under different growth conditions. Cell size quantified after five days of growth. Genotype and media treatment is indicated on the x-axis. Media type is indicated with either a white box for no treatment, or an orange box for media with 15µM thiamine and 500µM NAA. Mean and st. dev of distributions in pixels^2^: *mst2Δ epe1^deg^ sty1Δ* no treatment (123.0 ± 61.9), thiamine and NAA (39.1 ± 43.2); *mst2Δ epe1^deg^*no treatment (246.2 ± 74.6), thiamine and NAA (109.4 ± 64.0) (F) Percentage of *mst2Δ epe1^deg^ sty1+* and *mst2Δ epe1^deg^ sty1Δ* cells that survive following treatment with 15µM thiamine and 500µM NAA. Total colony count ratios were calculated by total number of colonies with thiamine and NAA divided by total number of colonies grown without thiamine and NAA. (G) *clr4+* mRNA expression measured by qRT-PCR in *mst2Δ epe1^deg^ sty1+* and *mst2Δ epe1^deg^ sty1Δ* after five days of treatment with 15µM thiamine and 500µM NAA. Log2 fold-change expression of mRNA is relative to expression without thiamine and NAA.

Mapping time-dependent changes across these GO categories reveals that the differential expression of cell proliferation and stress response genes subsides as adaptive Clr4 silencing is established (**Figure 4B, S4B-F**). Considering this apparent relationship, we wanted to interrogate the role that the stress response pathway plays in cell survival during heterochromatin misregulation and adaptive Clr4 silencing.

The CESR pathway plays a major role in *S.pombe* stress response. To interrogate the functional role of CESR during induced heterochromatin misregulation, we deleted the MAP kinase Sty1 in an *mst2Δ epe1^deg^* background. Sty1 regulates stress response in *S.pombe* by phosphorylating transcription factors that activate the expression of stress response genes, including a majority of genes in CESR (**Figure 4C**) ^58^. In our original *mst2Δ epe1^deg^*plate assay, when equal numbers of cells were plated, colony numbers were approximately equivalent regardless of Epe1 expression, suggesting a high heterochromatin misregulation stress survival rate (**Figure 4D**). To test how stress response plays into this survival, we plated *mst2Δ epe1^deg^ sty1Δ* cells on solid media for five days and measured cell colony size and survival frequency. *mst2Δ epe1^deg^ sty1Δ* colonies were on average smaller than *mst2Δ epe1^deg^* cells, both pre- and post-Epe1 depletion (**Figure 4E**). We observed only half as many colonies formed upon plating *mst2Δ epe1^deg^ sty1Δ* cells on NAA and thiamine-containing medium compared to *mst2Δ epe1^deg^* cells (**Figure 4F**). However, despite lower rates of stress-survival, Epe1-depleted *mst2Δ epe1^deg^ sty1Δ* colonies showed equally strong adaptive silencing of Clr4 transcription compared to *mst2Δ epe1^deg^* (**Figure 4G**). This suggests that the activation of stress response pathways is an on-pathway intermediate prior to adaptation, instead of directly driving redistribution of H3K9 methylation.^60^ Altogether, these results support Sty1 activity as beneficial for survival during heterochromatin misregulation.

### Loss of the RNA binding protein Red1 attenuates stress and delays adaptive *clr4+* silencing

We hypothesized that cells must leverage existing heterochromatin nucleation pathways to establish adaptive heterochromatin at new locations in the genome. This in turn could affect the duration and outcome of the stress phase and the subsequent adaptive phase. Based on our H3K9me3 ChIP-seq analysis of heterochromatin islands at meiotic genes and ncRNA, we focused on two major heterochromatin nucleation pathways – RNAi (Ago1, Dcr1) and MTREC (Red1).^46,56,61,62^ Surprisingly, we observed a lesser degree of *clr4+* transcriptional silencing in the adapted phase only in *red1Δ* cells but not *ago1Δ* or *dcr1Δ* cells (**Figure 5A, S5A**).^47^ To determine if the changes we measured in *clr4+* silencing correlate with any change in stress cells experience, we conducted growth experiments using replicate *mst2Δ epe1^deg^ red1Δ* cultures on plates supplemented with NAA and thiamine, as well as in continuous culture using the eVOLVER system (**Figure 5B, S5B-C**). These approaches confirmed a fitness increase during the stress phase compared to *red1+* cells. In contrast, *ago1*Δ and *dcr1*Δ cells exhibited the expected loss of fitness further confirming a distinctive role for Red1 during the stress and adaptive phase. To further quantify and compare loss of fitness during the stress phase, we calculated the mean minimum decrease in growth rates for each eVOLVER experiment (**Methods**). Red1-deficient cultures displayed a significantly smaller decrease in growth rate, similar to *mst2Δ epe1^deg^ clr4Δ*, compared to *mst2Δ epe1^deg^* or RNAi deletions (**Figure 5B**). We also compared Clr4 mRNA between untreated *red1Δ* and *red1+* and that this lesser degree of *clr4+* silencing was not due to pre-adaptation in untreated *red1Δ* (**Figure S5D**). Hence, these results suggest that MTREC mediated Clr4 recruitment may nucleate aberrant heterochromatin during the stress phase to drive downstream adaptation.^63–65^

**Figure 5:**
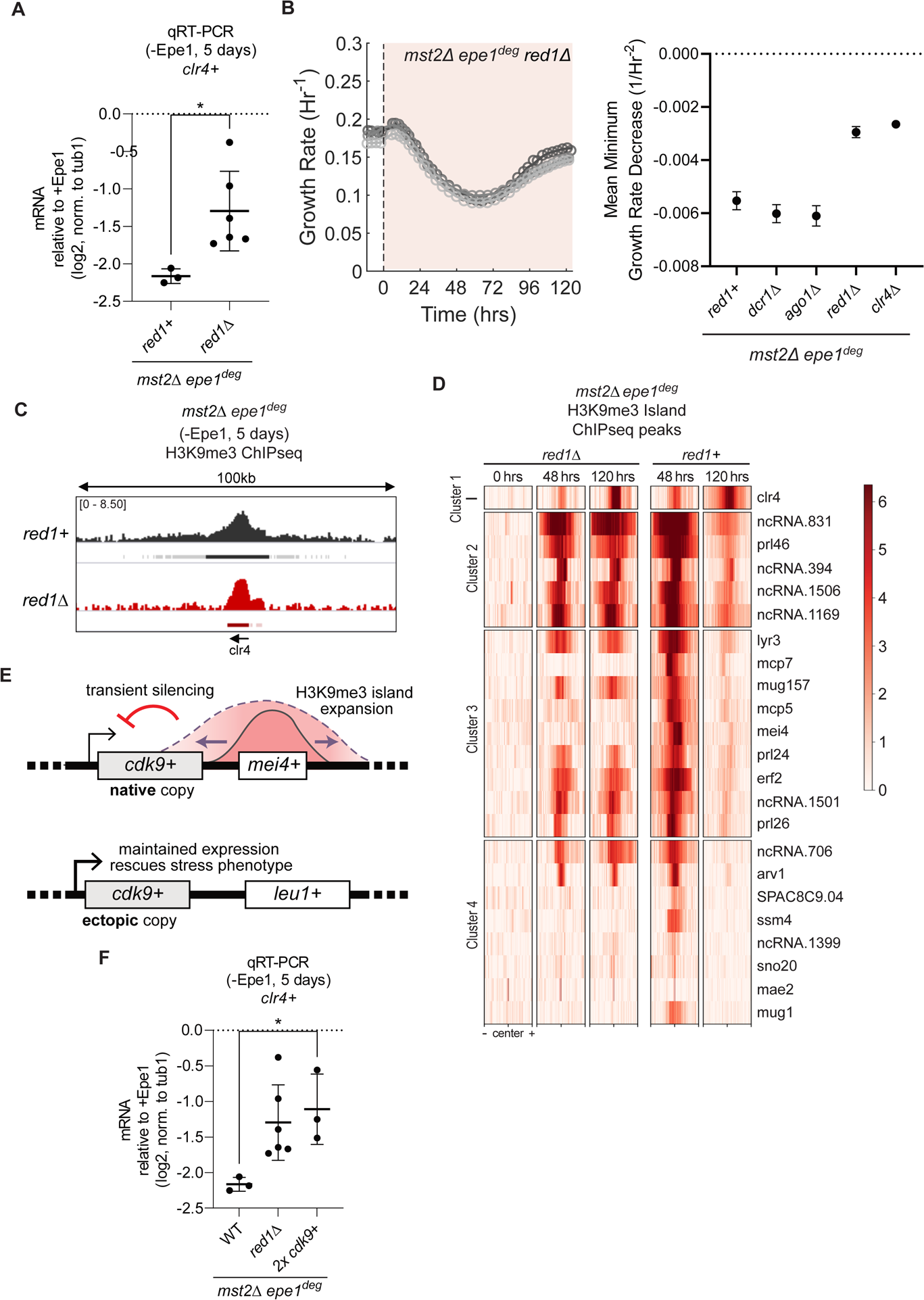
Loss of the RNA binding protein Red1 attenuates stress and delays adaptive clr4+ silencing. (A) *clr4+* mRNA expression measured by qRT-PCR after five days of treatment with 15µM thiamine and 500µM NAA. Log2 fold-change expression of mRNA is relative to mRNA expression without thiamine and NAA. Error bars represent standard deviation, N=3 or 6. Asterisk indicates p < 0.05. (B) (Left) Real-time monitoring of population growth rates of *mst2Δ epe1^deg^ red1Δ* cells cultured in eVOLVER. Treatment with 15µM thiamine and 500µM NAA was initiated at t=0hrs. Individual trendlines indicate replicates (N=2). Orange shaded portion represents the time period during which Epe1 has been depleted. (Right) Plot showing mean minimum decrease in growth rate for eVOLVER experiments of the indicated genotypes. N=3. (C) H3K9me3 ChIP-seq tracks of a 100kb window centered at the *clr4+* gene locus after five days of Epe1 depletion in *mst2Δ epe1^deg^ red1Δ* cells. Enrichment is shown in log2 fold change of IP normalized to input. Genotype is indicated on the left side of each track. Peaks identified are denoted in red below each track. The *clr4+* gene locus is specifically highlighted. (D) Heatmap showing loci of clustered H3K9me3 islands, originally identified in *mst2Δ epe1^deg^*, in *mst2Δ epe1^deg^ red1Δ*. Peaks are centered in a 24kb window and are clustered by K-means clustering from figure 3E. (E) Schematic for adding an ectopic copy of *cdk9+* at the *leu1+* locus. Heterochromatin misregulation allows spreading of the *mei4+* heterochromatin island over neighboring *cdk9+*. Insertion of *cdk9+* at the euchromatic *leu1+* locus prevents transient silencing from heterochromatin spreading. (F) *clr4+* mRNA expression measured by qRT-PCR after five days of treatment with 15µM thiamine and 500µM NAA. Log2 fold-change expression of mRNA is relative to mRNA expression without thiamine and NAA. Error bars represent standard deviation, N=3 or 6. Asterisk indicates p < 0.05.

To determine how heterochromatin changes in *mst2Δ epe1^deg^ red1Δ* cells lead to reduced stress and delayed *clr4+* silencing, we acquired batch cultures over a 120 hour period. ChIP-seq revealed the accumulation of H3K9me3 over the *clr4+* locus exhibited a significant contraction relative to *mst2Δ epe1^deg^ red1+* cells (**Figure 5C, S5E**). Additionally, during the stress period, *mst2Δ epe1^deg^ red1Δ* cells lost several H3K9me3 islands at meiotic genes, including *mcp7, mcp5, mei4, ssm4*, and *mug1,* consistent with a role for Red1 in nucleating these islands in cycling cells (**Figure 5D**).^56^ While the other H3K9me3 islands, originally identified in *mst2Δ epe1^deg^*, were still present, we observed that these remaining islands in *red1Δ* have less H3K9me3 enrichment at 48 hours compared to *red1+*. These H3K9me3 peaks also show less decay by 120 hours, possibly due to weaker *clr4+* silencing. These observations led us to test if any specific Red1 dependent H3K9me island expansions were primary drivers of the stress phase. We noted an expansion of H3K9me3 from the *mei4+* locus to a proximal gene, *cdk9+*. Cdk9 is an essential kinase that regulates various aspects of RNA polymerase II transcription including initiation, elongation and termination.^66–68^ Specifically, *cdk9+* is silenced during the stress phase (24-48 hours) but is derepressed once adaptation is complete (120 hours) (**Figure 3E,G**). To compensate for Red1 mediated *cdk9+* silencing, we inserted a second copy of *cdk9+* at the *leu1+* locus in *mst2Δ epe1^deg^* (*“2x cdk9+”*) (**Figure 5E**). Our rationale was that the second copy of *cdk9+* would not be subject to the transient silencing effects, enabling us to isolate the effects of silencing *cdk9+* from the general effects of depleting Epe1. Indeed, we observed a weaker growth defect in colony size upon depletion of Epe1, compared to the original strain with one copy of *cdk9+* (**Figure S5F**). This was complemented with reduced *clr4+* silencing to a level that mirrored *mst2Δ epe1^deg^ red1Δ,* suggesting that silencing of *cdk9+* is a key downstream event of Epe1 depletion for both adaptive *clr4+* silencing as well as the intermediate, low fitness stress phase (**Figure 5F**). Taken together, these results show that the expansion of heterochromatin islands, following the loss of Epe1, is crucial for cell stress and subsequent adaptation.

### Adaptive heterochromatin exhibits memory upon re-induction of stress

As we previously showed, depletion of Epe1 is rapid and efficient, occurring in as little as 30 minutes after the addition of NAA and thiamine (**Figure 1C**). Therefore, the *epe1^deg^* allele enables us to rapidly and reversibly cycle between Epe1 depletion and expression. To test whether cells that had adapted to Epe1 loss also exhibited memory, we restored Epe1 expression in adapted cells for different recovery periods. We refer to these recovery periods as **short** (24 hours), **medium** (48 hours) and **long** (72 hours). Following the recovery period, we re-initiated Epe1 depletion to generate a second stress phase (**Figure 6A-B**). If Clr4 silencing is faster during the second stress phase, it would imply that cells have the potential for adaptive memory, where an original lost adaptation can be recalled more quickly than the initial adaptive development.

**Figure 6.**
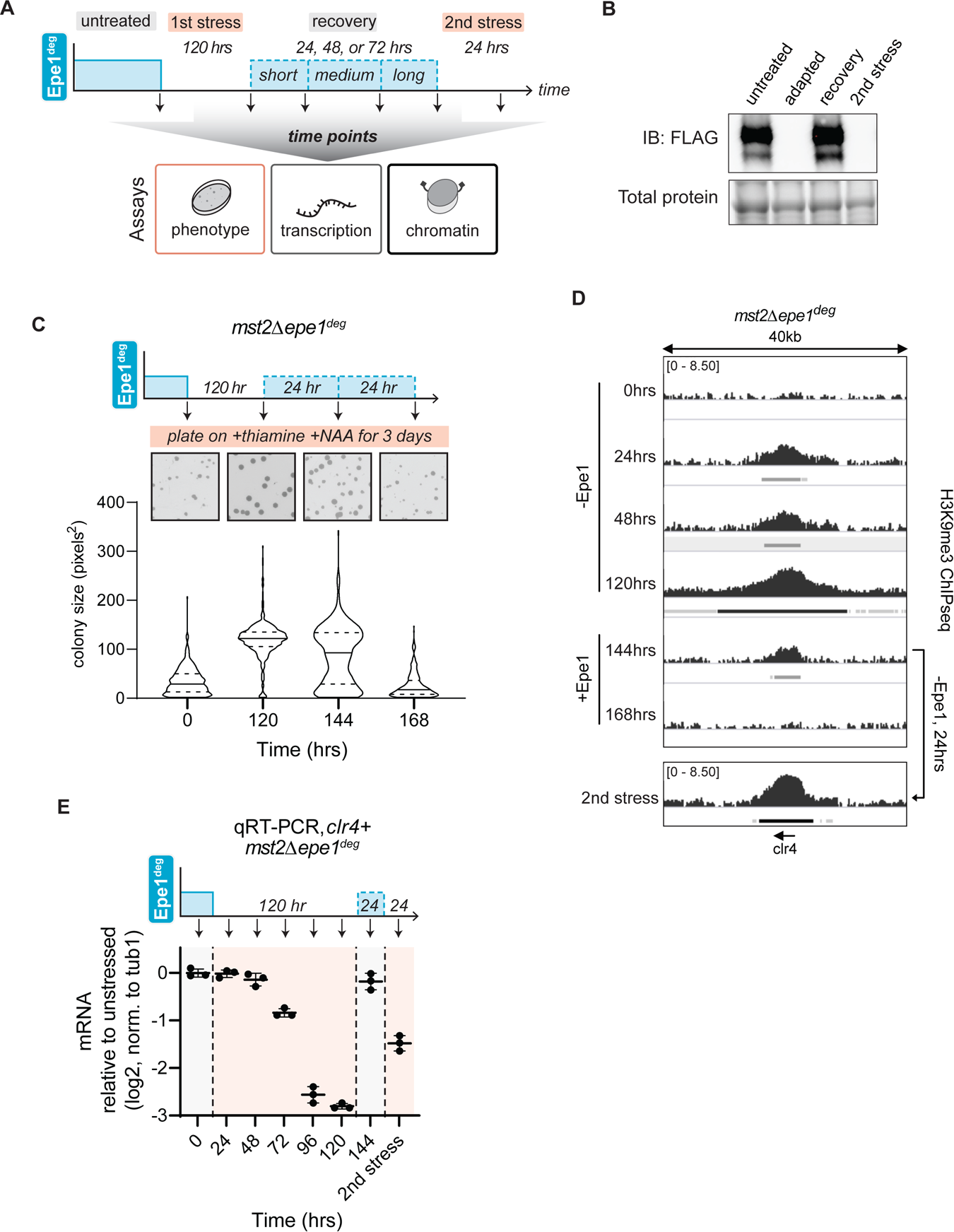
Adaptive heterochromatin exhibits memory upon re-induction of stress. (A) Schematic indicating cycling of Epe1 availability in *mst2Δ epe1^deg^*to measure epigenetic memory at the phenotype, transcription and chromatin level. Untreated cells express Epe1 and have not experienced heterochromatin misregulation. Adapted cells have experienced Epe1 depletion and silenced *clr4+* after 120 hours. Cells were allowed to recover with Epe1 expressed for 24, 48 or 72 hours, followed by a second round of Epe1 depletion. (B) Western blot for Epe1-3xFLAG-AID in *mst2Δ epe1^deg^* cells. Adapted and re-induced adapted cells were grown with 15µM thiamine and 500µM NAA for a minimum of 24hrs. Total protein levels are shown in the lower panel. (C) Colony size distribution of *mst2Δ epe1^deg^* cells, in pixel area, after three days of growth with Epe1 depletion from 15µM thiamine and 500µM NAA (Bottom). Epe1 expression and snapshots of culture plates (Top). Mean and st. dev of distributions in pixels^2^: 0 hours (34.5 ± 27.3), 120 hours (118.9 ± 37.6), 144 hours (87.3 ± 59.4), 168 hours (26.3 ± 25.9) (D) H3K9me3 ChIP-seq tracks in a 40kb window centered at the *clr4+* gene locus after five days of Epe1 depletion in an *mst2Δ* background. Enrichment is shown in log2 fold change of IP normalized to input. Epe1 was depleted within the window of 0-120 hours. After 120 hours, Epe1 was expressed for a recovery period of 24, 48, or 72 hours. Recovered populations were then put under a second stress by depleting Epe1 for another 24 hours.. Peaks identified are denoted below each track. The *clr4+* gene locus is specifically highlighted. (E) mRNA expression in *mst2Δ epe1^deg^* cells measured by qRT-PCR for *clr4+*. Orange shaded portion represents the time period during which Epe1 has been depleted. Epe1 expression and time points are shown above the figure.

As controls, untreated *mst2Δ epe1^deg^* cells exhibited smaller sized colonies with substantial heterogeneity upon Epe1 depletion (**Figure 6C**). Adapted cells formed uniformly sized colonies upon sustained Epe1 depletion. In contrast, adapted cells that had experienced a short recovery dose of Epe1 (Epe1 re-expressed for 24 hours) produced an intermediate colony size phenotype, with a bimodal distribution of small and large colonies. These results suggest that some proportion of short recovery cells had reverted to the untreated state while others preserved the adapted state during the 2^nd^ stress phase. Medium and long recovery cell populations produced colony size phenotypes that matched untreated cells, suggesting that a prolonged recovery phase (>24 hours) led to the complete loss of adaptive memory. Hence, our results reveal that a memory associated with heterochromatin misregulation can persist for about 24 hours (∼6-8 cell generations) following the removal of the initiating stress.

To measure memory at the chromatin level, we performed H3K9me2 and H3K9me3 ChIP-seq on recovering *mst2Δ epe1^deg^* cells. A short recovery over 24 hours led to a reduction of H3K9me2 and H3K9me3 (equivalent to 24 hours of stress), and by medium recovery, adaptive H3K9me2 and H3K9me3 at Clr4 had decayed to undetectable levels (**Figure 6D, S6**). To test if cells in short recovery could reestablish silencing at the *clr4+* locus, we reintroduced heterochromatin stress to short recovery cells by depleting Epe1 a second time. After stress reintroduction, short recovery cells re-established H3K9me3 much faster than untreated cells (24 hours in short recovery cells versus 72 hours in naive cells) (**Figure 6D**). These changes in Clr4 adaptive heterochromatin are mirrored in *clr4+* transcription, as well. Recovering cells expressed *clr4+* at unstressed levels, while cells that have had a prior experience of stress quickly reestablished *clr4+* adaptive silencing (**Figure 6E**). These results show that novel adaptive H3K9 methylation is maintained for several cell divisions during stress recovery, and this residual methylation can encode epigenetic memory to more rapidly reestablish adaptive silencing.

### The time scale of adaptive memory can be tuned by histone acetylation

Cells may need to extend or even erase memory associated with past events.^33,36^ To identify other chromatin-based mechanisms that might modulate and enhance adaptive memory duration, we considered the interdependence of H3K9me with other histone modifications. Histone hyperacetylation (H3K9Ac, H3K14Ac, H3K18Ac) and histone turnover are characteristic features of actively transcribed genes, and loss of histone acetylation can promote heterochromatin inheritance.^69,70^ Although Mst2 has been deleted in our strains (*mst2Δ epe1^deg^*), it is possible that other histone acetyltransferases could play additive roles in tuning the length of adaptive memory.^71^ We deleted a second acetyltransferase, Gcn5, in a *mst2Δ epe1^deg^* background (*mst2Δ gcn5Δ epe1^deg^*). Cells with the *mst2Δ gcn5Δ epe1^deg^*genotype grew comparably to *mst2Δ epe1^deg^* prior to Epe1 depletion, and indicated a slightly stronger stress phenotype after loss of Epe1 (**Figure S7A**). Using this strain, we performed memory experiments as previously described where we now cycled *mst2Δ gcn5Δ epe1^deg^* cells between conditions where Epe1 was depleted or expressed. Unlike what we observed in cells where Gcn5 is present (memory decayed after the short recovery period), *mst2Δ gcn5Δ epe1^deg^* cells retained memory after short, medium, and long recovery periods. After each recovery period, colony sizes most closely resembled adapted colonies, implying that these cells may exhibit prolonged, adaptive memory (**Figure 7A**).

**Figure 7.**
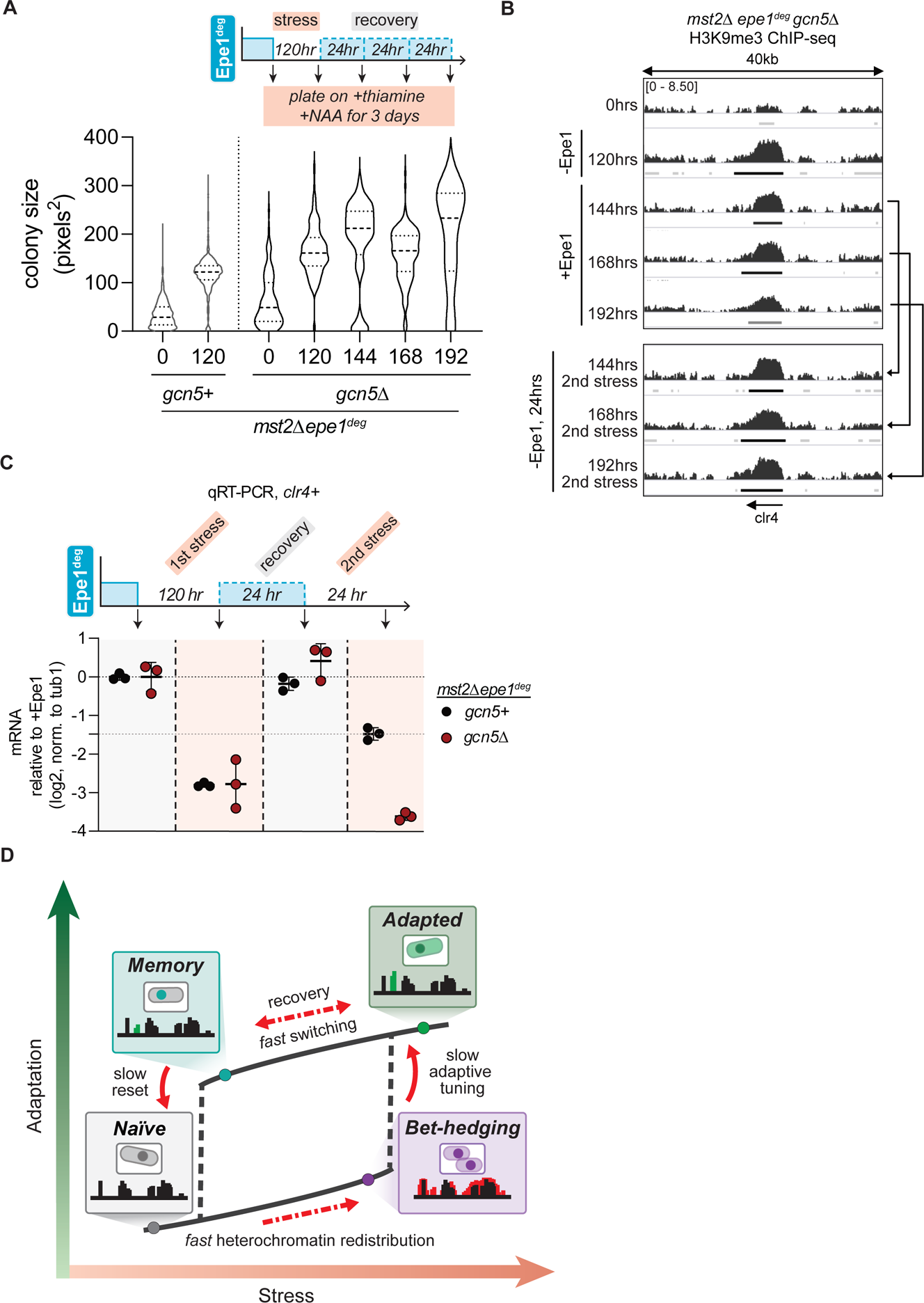
The time scale of adaptive memory can tuned by histone acetylation. (A) Colony size distribution of *mst2Δ epe1^deg^ gcn5Δ* cells, in pixel area, after three days of growth with Epe1 depletion from 15µM thiamine and 500µM NAA. Distributions of *mst2Δ epe1^deg^* colony sizes are shown for comparison. Mean and st. dev of distributions in pixels^2^: 0 hours (61.9 ± 53.8), 120 hours (166.1 ± 50.9), 144 hours (185.8 ± 87.9), 168 hours (158.2 ± 54.0), 192 hours (201.1 ± 102.3) (B) H3K9me3 ChIP-seq tracks in a 40kb window centered at the *clr4+* gene locus after five days of Epe1 depletion in an *mst2Δ epe1^deg^ gcn5Δ* background. Enrichment is shown in log2 fold change of IP normalized to input. Epe1 was depleted within the window of 0-120 hours. After 120 hours, Epe1 was expressed for a recovery period of 24, 48, or 72 hours. Recovered populations were then put under a second stress by depleting Epe1 for another 24 hours. Peaks identified are denoted below each track. The *clr4+* gene locus is specifically highlighted. (C) *clr4+* mRNA expression in *mst2Δ epe1^deg^ gcn5+* and *mst2Δ epe1^deg^ gcn5Δ* cells measured by qRT-PCR. Orange shaded portion represents the time period during which Epe1 has been depleted. (D) Heterochromatin-defining H3K9 methylation (H3K9me), normally reserved to maintain genome integrity can be redistributed across the genome to establish new and potentially adaptive phenotypes. Establishing adaptive H3K9me patterns occurs over surprisingly slow time-scales relative to the initiating stress. The slow kinetics of adaptation may serve as a bet-hedging strategy for cells to decipher optimal survival solutions. Upon removal of stress, cells relax to new transcriptional and chromatin states rather than revert to their initial (ground) state, establishing a tunable memory for a future adaptive epigenetic response. Cells exhibit history dependence wherein a prior exposure to stress locks cells in a new transcriptional state that encodes adaptive memory.

Next, we measured other molecular correlates of memory-*clr4+* mRNA levels and H3K9me3 levels at the *clr4+* locus using ChIP qPCR in *mst2Δ epe1^deg^* and *mst2Δ epe1^deg^ gcn5Δ* populations. We made two important observations: First, during each recovery phase, *clr4+* mRNA levels reverted to high levels of transcription, matching what we typically observed in untreated cells. Supporting our model that H3K9me3 is required for epigenetic adaptive memory, all recovery phase *mst2Δ epe1^deg^ gcn5Δ* cells retained H3K9me2 and H3K9me3 at the *clr4+* locus (**Figure 7B, S7B**). Hence, altering the chromatin state and the persistence of H3K9me3 at the *clr4+* locus has an additive effect on adaptive memory. By measuring total H3K14ac through western blotting, we found that deleting Mst2 led to a reduction in H3K14Ac and additionally deleting Gcn5 in this background eliminated nearly all available H3K14Ac in the cell population (**Figure S7C**). These results suggest that enhanced H3K9me, which arises from the absence of H3K14Ac, can tune the length of adaptive memory. This interdependence on post-translational modifications may allow for cells to rapidly toggle adaptive silencing states enabling them to extend (or erase) memory of past stress events.

Second, although H3K9me3 levels are higher in *gcn5Δ* than in *gcn5+* cells, the modification itself was not sufficient to have a repressive function during the recovery phase. Instead, *clr4+* adaptive silencing returned in an Epe1 depletion-dependent manner. When recovering *mst2Δ epe1^deg^ gcn5Δ* was exposed to a 2^nd^ stress, cells re-established *clr4+* silencing even though adaptive H3K9me3 maintained a similar distribution compared to the recovering population (**Figure 7B-C**). Taken together, our results imply that while H3K9 methylation has an important role in preserving memory associated with Clr4 silencing, it may not be the only factor that contributes to this process (**Figure 6D-E, S6, 7B-C**). Therefore, we tested whether short recovery cells exhibited unique transcriptional changes that could account for adaptive memory and silencing by performing RNA-seq analysis on short and medium recovery *mst2Δ epe1^deg^* cell populations. Indeed, PCA reveals that cells during recovery phases exhibit unique transcriptional dynamics that were distinct from untreated, stress, or adapted phase cells (**Figure S7D**). Importantly, the cell transcriptional state when Epe1 is re-expressed does not instantaneously revert to the initial untreated state. Transcriptional changes in the short recovery phase were primarily enriched for genes related to ncRNA and rRNA processing, ribosome biogenesis, and RNA metabolic processes (**Figure S7E**). We also observed enrichment for CESR transcripts, suggesting that the shift from the adapted state and loss of heterochromatin at *clr4+* activates part of a stress response. Since we have previously implicated MTREC in adaptive Clr4 silencing, we examined known Red1 targets (**Figure S7F**).^73,74^ We observed large transcriptional changes in Red1 targets during both the stress and recovery phase, reinforcing the causal role that MTREC has in adaptive Clr4 silencing.

## DISCUSSION

Cells can repurpose and leverage existing epigenetic pathways to modulate gene expression states in response to environmental change. In fission yeast, H3K9 methylation exhibits adaptive epigenetic potential that is unlocked when cells encounter various environmental stressors, including anti-fungals, caffeine, or nutrient restriction.^21,39,57,75,76^ These stressors impact two major H3K9me regulators, Epe1 and Mst2 at the RNA and protein level. Our ability to control Epe1 and Mst2 levels (in our *mst2Δ epe1^deg^* and *mst2^deg^ epe1^deg^* strains), rapidly and reversibly enables us to mimic the natural stress response of fission yeast cells, independent of an initiating stress. Based on previous research and our work, we propose the following model for epigenetic adaptation. Exposure to stress impacts Epe1 and Mst2 levels in *S.pombe* cells. Epe1 undergoes proteosome mediated degradation while Mst2 produces a shorter, MYST-domain deficient isoform in a stress-dependent manner.^21,57^ Manifesting a stress-driven slow growth phenotype would allow time for an adaptive silencing pathway to establish H3K9me at new locations in the genome. While our inducible depletion system has several controlled variables that separate it from a natural system, it undeniably provides a window into the earliest transcriptomic changes that cells undergo prior to adaptation.

Our system enables the capture of early and intermediate transcriptional states that lead to adaptive silencing of *clr4+* which would be impossible to capture using conventional genetic methods.^47^ Notably, we found that the decision for cells to adapt and silence *clr4+* is a slow process, taking up to 120 hours, which is two orders of magnitude slower than the timescale of Epe1 depletion (approximately 30 minutes). We propose that these slow dynamics may represent a bet-hedging strategy, where cells reversibly sample different transcriptional states to enhance fitness before converging on an optimal solution.^77^ Our findings parallel stochastic tuning models in budding yeast, which posit that transcriptional noise is positively selected to promote cell survival in novel environmental conditions.^24,25^

By analyzing the transcriptomic profile of cells, we observed transient activation of stress response pathways correlating with a slow growth phenotype. Activation of genes associated with the core environmental stress response (CESR) serve as an on-pathway intermediate preceding adaptation.^60^ These dynamics strikingly resemble how cancer cells often vexingly develop resistance, resulting in poor prognosis and treatment outcomes. For example, glioblastoma cells transiently exit the cell cycle, exhibit slow growth, misregulate H3K27 methylation-dependent epigenetic pathways, and ultimately adapt by entering a state that is refractory to chemotherapeutic interventions.^19^ Bacteria also enter slow-growing persister states through the activation of a stress-induced SOS pathway, leading to genetic changes and antibiotic resistance.^8,78^ In both instances, slow growth is a common denominator that is triggered by the activation of the stress response pathway. Thus, the activation of stress response pathways may represent a general principle that cells leverage to explore adaptive phenotypes when exposed to novel environments.^60^

It is possible that the initial, unregulated expansion of H3K9me during early heterochromatin misregulation leads to the silencing of many essential genes that disrupt fitness and cell survival. We speculate that this could represent a temporary switch in growth strategy to a slow proliferation state until beneficial adaptations can be acquired. The expansion of heterochromatin domains over essential genes could also act as a filter for stress and adaptive responses. Since the spatial expansion of heterochromatin must build over time, cells would need to experience sustained exposure to a stressor before committing to adaptive H3K9me redistribution, preventing premature adaptive responses to transient environmental perturbations.

Establishing adaptive heterochromatin at the *clr4+* locus follows unique dynamics that are distinct from H3K9me-spreading at other regions during heterochromatin misregulation. These dynamics suggest active recruitment of heterochromatin initiation and maintenance factors that have adaptive potential. This process is partially dependent on Red1, as *mst2Δ epe1^deg^ red1Δ* cells consistently show weaker *clr4+* adaptive silencing within our 120-hour window, and an increase in fitness during the stress phase. This raises the possibility that adaptive silencing is mediated by co-transcriptional or post-transcriptional processes that involve non-coding RNA recognition by factors that are part of the MTREC complex.^56,61,62^ The expansion of Red1-dependent H3K9me islands during the stress phase contributes to activating stress response pathways, thus explaining why deleting Red1 alleviates stress and leads to slower *clr4+* silencing. How this RNA elimination machinery affects the formation of *de novo* adaptive heterochromatin in other stress contexts, and the extent to which it can be repurposed when cells encounter novel environments, requires further investigation. Our results from relocating *cdk9+,* an essential gene that is subject to transient Red1 silencing during the stress phase, support an essential role for Red1 during stress and subsequent adaptation. These results also reveal how the arrangement of genes on chromosomes could confer unforeseen advantages for cell survival when exposed to acute stress. This is particularly intriguing given that the *mei4+* and *cdk9+* preserve synteny even in the highly diverged *Schizosaccharomyces japonicus,* suggesting how the potential for organisms to adapt could be an emergent property that shapes genome organization.^79^ Additionally, our results mirror recent work where inhibition of the human CDK9 ortholog produces transcriptional reprogramming, supporting a model where the inhibition or transient repression of essential genes encourages epigenetic adaptations.^80^ How other transiently-silenced essential genes, including the previously identified *cup1+* gene, modulates the formation of epigenetic adaptations requires further investigation.

Our unique ability to toggle between Epe1 removal and expression enabled us to determine that cells can retain a memory associated with prior heterochromatin misregulation for approximately 6-8 generations following the initial stress. This memory depends on residual H3K9 methylation which qualitatively resembles earlier work where adding back Epe1 permanently poises cells in a novel, fixed epigenetic state.^81^ In contrast, re-expressing Epe1 in our inducible system leads to total *clr4+* derepression despite significant H3K9me3 being present during the recovery phase. Retained H3K9me3 levels at the *clr4+* locus are even more strongly retained in recovering *mst2Δ gcn5Δ epe1^deg^* cells, and yet *clr4+* continues to be expressed upon the reintroduction of Epe1. One possibility is that H3K9me3 is required for memory but silencing requires additional inputs from other factors involved in different tiers of transcriptional, co-transcriptional and post-transcriptional regulation.^82–85^ We speculate that memory and subsequent adaptive silencing may depend, not only on H3K9me3, but also on the cell being poised to adapt due to novel network-level gene expression changes.^27,54,86^ Our RNA-seq measurements and PCA analysis firmly support the idea of new adaptive cell states that can potentially encode memory. Cells in which Epe1 has been reintroduced during the recovery phase exhibit unique gene expression signatures, distinct from the untreated state. We propose that this feature represents a form of cellular hysteresis, or history dependence, in which case the gene regulatory network in cells is completely rewired upon either removing or adding back Epe1 (**Figure 7E**).^87^ It remains to be seen whether other epigenetic regulators also exhibit hysteresis given the slow kinetics of establishing novel epigenetic states *de novo*.^88^

In conclusion, our inducible system uniquely allowed us to capture the highly dynamic and unexpected changes in gene expression and chromatin states following acute heterochromatin misregulation in yeast populations, changes that would be otherwise obscured in conventional genetic assays. Using this approach, we could faithfully reconstruct the pathways that cells undertake prior to and during adaptation as well as investigate how the adapted state is memorized across multiple generations. Our findings reveal several key and distinguishing features of adaptation by epigenetic responses, including the slow kinetics of the process consistent with the concept of epigenetic bet-hedging, and the establishment of adaptive memory that is influenced by cellular history. This work also illuminates strategies by which cells stabilize new gene expression programs to endure environmental change, with implications across biology from development to evolution. Ultimately, we demonstrate a powerful experimental framework to probe adaptation mediated through chromatin regulation - an exciting frontier offering new insights into phenotypic plasticity and organismal responses to transient stresses.

## LIMITATIONS OF THE STUDY

Our work utilizes an inducible, on-demand system to initiate and track an adaptive epigenetic response upon removing or adding back key heterochromatin regulators in *S.pombe*. Natural stressors would likely elicit additional interspersed transcriptomic changes beyond what we observed. For example, caffeine affects DNA replication, so cells would experience both an adaptive response and replication stress. Nevertheless, understanding how entangled transcriptomic changes enable successful adaptation to stress will be an important area for future investigation. Our current population-level studies do not capture cell-to-cell heterogeneity in the specific choice of *clr4+* as the primary locus enabling the observed epigenetic adaptation. By profiling transcriptomic changes at the single cell level over time, future work could delineate the diversity of molecular paths individual cells take before converging on an apparent optimal solution. These experiments may reveal whether the adaptive mechanism is purely stochastic across cells or if certain deterministic factors target the response to specific loci. Capturing single cell dynamics will add a valuable layer of understanding on top of the population-wide measurements we have conducted so far.

## METHODS

### Yeast strains, plasmid construction, and culturing

All *S. pombe* yeast strains used in this study were generated using either established methods for lithium acetate or electroporation transformations, or by meiotic crossing followed by tetrad dissection. All strains were genotyped using a colony PCR protocol. Plasmid constructs to create modified nmt81-Epe1-3xFlag-AID and nmt81-Mst2-3xFlag-AID inserts were constructed by modifying an existing pFA6a 3xflag AID IAA-17 degron kanMX6 plasmid. Full plasmids were made using ligation methods following PacI digestion to insert the nmt81 promoter. This insert repaired the PacI site, allowing for a second PacI digestion to insert the CDS for Epe1 or Mst2.

EMMC was used as the base media for all cell culturing experiments, and all cultures were grown at 32C. For experiments involving sampling cultures over a time-course, a small volume of cells from each timepoint culture was used to nucleate the culture for the next timepoint in the appropriate media. All strains used in this study are listed in Table 1.

For colony size quantification, seed cultures were grown overnight at 32C in EMMC in liquid media. Seed cultures were then used to nucleate fresh liquid cultures at a low starting OD (< 0.3 OD) and allowed to grow for about 6-8 hours. An equivalent number of cells were then diluted and plated on solid media, either EMMC or EMMC media supplemented with 15µM thiamine and 500µM NAA and spread with sterile glass beads. Plated cultures were grown at 32C, and pictures were taken at 3 and 5 days from plating on a Biorad ChemiDoc with white epifluorescence. Images were analyzed in FIJI for trimming plate edges, identifying individual cell colonies, and quantifying colony number and size. To calculate percentage survival, we calculated colony count ratios between EMMC and EMMC media supplemented with 15µM thiamine and 500µM NAA.

### Western blotting

To test time-dependent depletion of Epe1, cultures were seeded at a low OD (∼0.3) in liquid media EMMC or EMMC supplemented with 15µM thiamine and 500µM NAA. For later memory experiments that switched Epe1 expression, cells cultured for five days in 15µM thiamine and 500µM NAA were harvested and a portion of the culture was used to start a new overnight culture in EMMC. This EMMC culture was then both harvested after 24 hours and used to inoculate a new culture in 15µM thiamine and 500µM NAA for a second time. That culture was then sampled after an additions 24 hours. All cultures were harvested by centrifuging 3-5 OD, decanting supernatant, and storing harvested pellets at -80C.

To extract protein for immunoblotting, cell pellets were processed using a standard TCA precipitation protocol. Pellets were washed with 1mL of ice cold water, then resuspended in 150uL of YEX buffer (1.85 M NaOH, 7.5% beta-mercaptoethanol). Resuspended pelleted were incubated on ice for ten minutes, then 150uL of 50% TCA was mixed into each sample and incubated for ten minutes on ice. Samples were then centrifuged for 5 minutes at 13000 rpm at 4C, after which the supernatant was decanted. Pellets were then resuspended in SDS sample buffer (125mM TRB pH 6.8, 8M urea, 5% SDS, 20% glycerol, 5% BME) and centrifuged for 5 minutes at 13000 rpm at 4C. Samples were then run on an SDS page gel at 45 minutes at 200V. Stain-free imaging was performed on a Biorad ChemiDoc. Gel transfer was then performed on a Trans-Blot Turbo Transfer to a nitrocellulose membrane. Immunoblotting was performed by blocking the nitrocellulose membrane with 5% non-fat dry milk in Tris-buffered saline pH 7.5 with 0.1% Tween-20 (TBST) for about an hour. Blots were then incubated overnight with primary antibody at 4C, then washed with TBST three times and incubated with secondary antibody for an hour. Incubated blots were imaged using enhanced chemiluminescence on a Biorad ChemiDoc.

### eVOLVER growth assay

Continuous culture experiments were performed in eVOLVER, designed and set up as previously described ^52^. Two replicate cultures of each strain were grown in 25 mL of EMMC media at 32 C. Growth was maintained in log phase using “turbidostat” mode to constrain optical density between 0.1 and 0.6. When cultures rise beyond the maximum OD, a dilution event is triggered, and growth rate is calculated for the duration since the previous dilution by fitting OD measurements to the exponential equation: *OD*600 = (*initial density*) ∗ *e*^(*growth rate*) ∗(*time*)^. Media condition changes were executed by spiking individual culture vials as well as the input EMMC media with a 1000x concentrated solution of thiamine + NAA in DMSO. Influx and efflux operations were manually triggered to flush untreated media from the lines.

To calculate the time derivative decrease in growth rate post-Epe1 depletion, first 60 hrs (2.5 days) after addition of thiamine and NAA were considered. Time derivatives of growth rate were calculated at each pair of consecutive growth rates with MATLAB’s gradient function. A moving average with a sliding window of length 3 was applied to the time derivative of the growth rate, and the minimum of this moving average was found to be the minimum change in growth rate for each experiment vial. Subsequently, the average and standard deviation of the minimum change in growth rate was calculated across triplicate experiment vials.

### qRT-PCR and RNA sequencing analysis

Cultures were grown in liquid culture containing either EMMC media or EMMC media supplemented with 15µM thiamine and 500µM NAA. For *mst2Δ epe1^deg^* time points 0-120hrs, cells were cultured and harvested from eVOLVER. For memory RNA experiments, *mst2Δ epe1^deg^* cells were grown in manually maintained incubated cultures. Cells were grown to 0.3-1.0 OD and harvested by centrifuging ∼10mL of culture at 2000 rpm for 2 minutes. Cell pellets were washed once in distilled water, centrifuged at 5000 rpm for 30 seconds, and stored at -80C.

Stored pellets were thawed on ice for 5 minutes, then resuspended in 750uL TES buffer (0.01M Tris pH7.5, 0.01M EDTA, 0.5% SDS). 750uL acidic phenol chloroform was immediately added afterwards, samples were vortexed, and then incubated on a heat block at 65C. Samples were incubated for a total of 40 minutes, with 20 seconds of vortexing every 10 minutes. Afterwards, heated samples were placed on ice for 1 minute, shaken, and transferred to phase lock tubes. Phase lock tubes were centrifuged for 5 minutes at 13,000 rpm at 4C, and the aqueous phase was transferred to a clean Eppendorf tube and ethanol precipitated. Isolated nucleic acids were then treated with DNAse I at 37C for 10 minutes and cleaned up on Qiagen RNeasy Clean-Up columns. Purified total RNA was converted to cDNA by annealing reverse primers complementary to target genes and reverse transcribing with SuperScript III Reverse Transcriptase (Invitrogen). qRT-PCR was performed with SYBR Green dye on a CFX Opus 384 Real-Time PCR System. All qRT experiments were reproduced for at least three independently growth replicates.

Libraries were prepared and sequencing was performed commercially. Raw fastq files were evaluated using FastQC (v0.11.9) and trimmed using Trimmomatic (v0.39) and aligned to the ASM294v2 *S. pombe* reference genome using STAR (v2.7.8a) then indexed using samtools (v1.10) (Andrews, 2010; Bolger et al., 2014; Dobin et al., 2013; Danecek et al., 2021). Bam files were grouped by genotype replicate and differential expression analysis was performed through Defined Region Differential Seq in the open source USEQ program suite (v9.2.9) (http://useq.sourceforge.net;Love et al., 2014). The cutoff for significant differential expression of pairwise gene comparisons was defined as a P value of <0.01 (prior to phred transformation) after Benjamini and Hochberg multiple testing corrections. For principal component analysis, rlog counts were used to perform MDS analysis, and custom ggplot2 R scripts were used to generate scatterplot figures. Volcano plots were drawn using the ggplot2 library, and heatmaps were drawn using the pheatmap library, as well as the standard R library and functions. Gene Ontology analysis was performed using the web-based tool AnGeLi with a p-value cutoff of < 0.01 with FDR correction for multiple testing and default settings (Britton et al., 2015). Raw and processed data are deposited in GEO under the accession number GSE235808.

### Chromatin immunoprecipitation, ChIP-seq library preparation and analysis

Cultures were grown in liquid culture containing either EMMC media or EMMC media supplemented with 15µM thiamine and 500µM NAA in manually maintained incubated cultures. Cells were grown to mid-log phase (0.9-1.6 OD) and then harvested by fixation with 1% formaldehyde for 15 minutes then quenched with glycine for 5 minutes. Fixed cultures were then centrifugated, washed twice with 1xTBS, and stored at -80C. To process samples, frozen pellets were thawed at RT for 5 minutes, then resuspended in 300 uL chip lysis buffer (50 mM HEPES-KOH, pH 7.5, 100 mM NaCl, 1 mM EDTA, 1% Triton X-100, 0.1% SDS, and protease inhibitors). Glass beads (500uL, 0.5mm) were added to each tube and cells were lysed by bead beating in an Omni Bead Ruptor at 3000 rpm × 30 s × 10 cycles. Ruptured cells were then collected by using a heated sterile needle to pierce the bottom of each tube, then collecting the lysate in a fresh tube via centrifugation. Lysate was then sonicated in a Q800R3 Sonicator to fragment sizes ranging from 100-500 base pairs. Sonicated lysate was then centrifuged at 13,000 rpm for 15 minutes at 4C, and the liquid portion was transferred to a new tube. Protein content was normalized using a Bradford assay. 25uL of each sample was reserved as input, to which 225uL 1xTE/1%SDS was added. Protein A Magnetic Dynabeads were preincubated with either Anti-H3K9me2 [Abcam, ab1220] or Anti-H3K9me3 [Active Motif, 39161] antibody. 30uL beads preincubated with 2ug antibody was added to 500uL cell lysate and incubated for 3 hours at 4C. Beads were held on a magnetic stand for subsequent washing cycles. For each wash cycle, cells were centrifuged at 1000 rpm for 1 minute at 4C, placed on the magnetic stand and allowed to settle, then liquid was removed by vacuum pipette. Then 1mL wash buffer was added and samples were rotated for 5 minutes per wash. Samples were washed three times with chip lysis buffer, then once with 1xTE. Samples were then eluted by suspending the beads in 100uL 1xTE/1%SDS for 5 minutes at 65C, then extracting liquid. A second elution was performed with 150uL 1xTE/0.67%SDS. Input and immunoprecipitated samples were then incubated overnight at 65C. We then added 60ug glycogen, 100ug proteinase K, 44uL of 5M LiCl, and 250uL of 1xTE was added to each sample and incubated at 55C for 1 hour. DNA was then extracted using phenol chloroform extraction, followed by ethanol precipitation. Ethanol precipitated pellets were resuspended in 100uL 10mM Tris pH 7.5 and 50mM NaCl. qPCR was performed with SYBR Green dye on a CFX Opus 384 Real-Time PCR System. All ChIP experiments were reproduced for at least two independently grown replicates.

Libraries were prepared following the standard protocol for the NEBNext Ultra II DNA Library Prep kit. Libraries of *mst2Δ epe1^deg^ gcn5Δ* cells were sequenced on an Illumina Miseq, and all other libraries were sequenced on an Illumina Nextseq instrument. Raw fastq reads were evaluated using FastQC (v0.11.9) and trimmed using Trimmomatic (v0.39) (Andrews, 2010; Bolger et al., 2014). Trimmed reads were aligned to the ASM294v2 *S.pombe* reference genome using the Burrows-Wheeler Aligner (v0.7.17) and bam files were further processed using samtools (v1.10) (Li and Durbin, 2010; Danecek et al. 2021). Bedgraph coverage files were generated using deepTools (v3.5.1) and normalized IP against input in SES mode (Ramírez et al., 2016; Diaz et al., 2012). ChIP-seq H3K9me3 peaks were called using MACS2 with -g 12.57e6 in broad mode with a cutoff of 0.05 (Zhang et al., 2008). Bedtools intersect (v2.27.1) was used to identify genes overlapping with identified peaks. Heatmaps were generated using deepTools (v3.5.1) (Ramírez et al., 2016). Specific peak histograms were generated using the SushiR package and custom R scripts. Raw and processed data are deposited in GEO under the accession number GSE235808.

## ACKNOWLEDGMENTS

The authors declare no competing interests. We thank Danesh Moazed for sharing fission yeast strains used in this study. We thank Nidhi Khurana and Gulzhan Raiymbek for their support in obtaining preliminary data during the initial phase of this study. We thank Basila Moochickal Assainar, Amanda Ames, and Sumanth Maheshwaram for their supportive and insightful feedback regarding this study. We thank Tommy V. Vo for helping with ChIP-seq analysis. This work was supported by the National Science Foundation (NSF) (grant no. EF-1921677 to K.R. and A.S.K.), National Institutes of Health (NIH) (grant nos. R35GM137832 to K.R.; R01EB029483, R01EB027793, and R01AI171100 to A.S.K.), American Cancer Society (grant no. RSG2211701DMC to K.R.); T32GM007544 to A.L.; T32GM007315 to M.S.), Department of Defense Vannevar Bush Faculty Fellowship (no. N00014-20-1-2825 to A.S.K.), and Schmidt Science Polymath Award (no. G-22-63292 to A.S.K.).

