## Supplementary material for "Mapping the dynamics of epigenetic adaptation during heterochromatin misregulation": Supp_figs_and_tables

Supp. 1

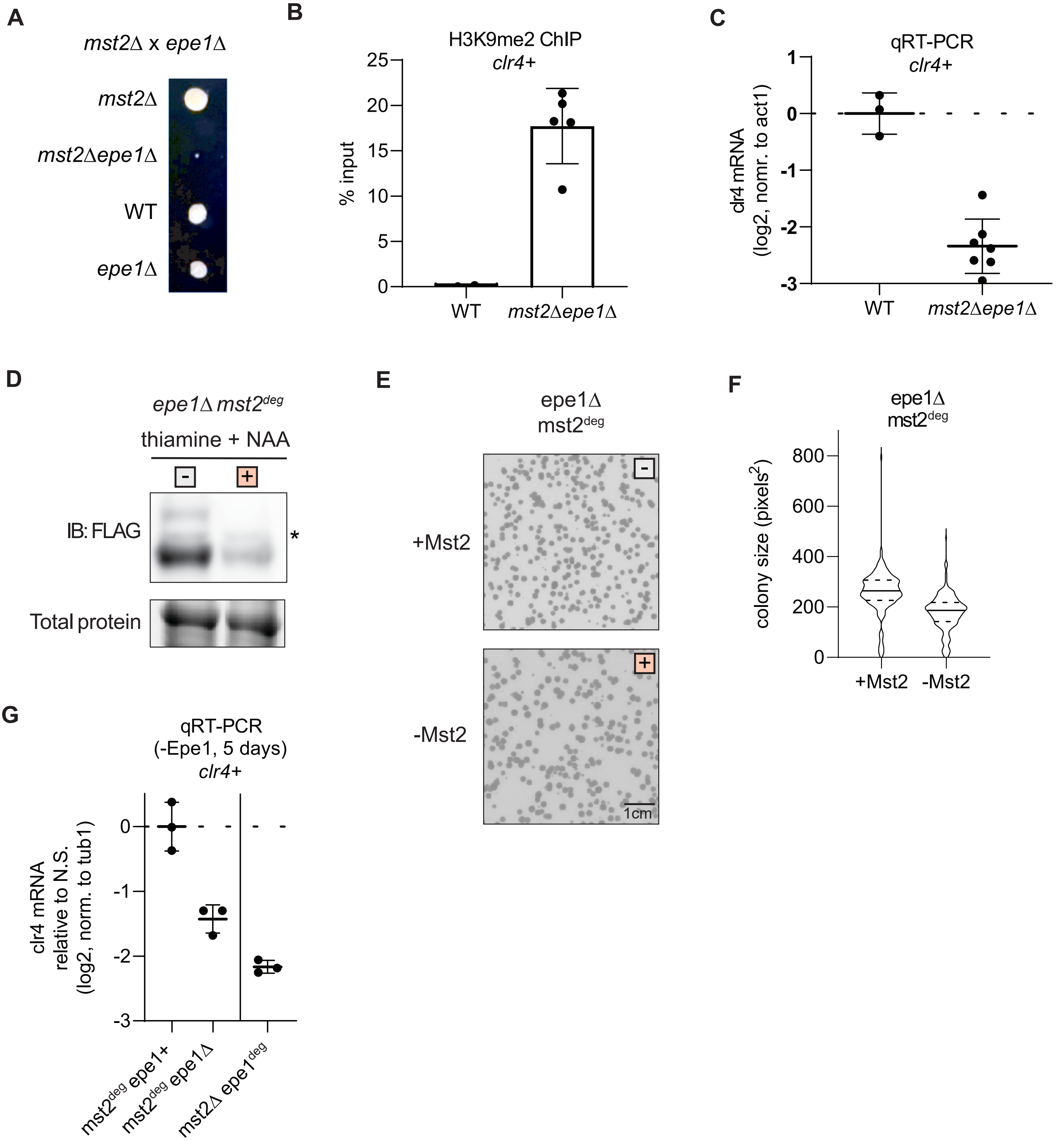

A
mst2Δ x epe1Δ
mst2Δ
mst2Δepe1Δ
WT
epe1Δ
B
H3K9me2 ChIP
clr4+
% input
25
20
15
10
5
0
WT
mst2Δepe1Δ
C
qRT-PCR
clr4+
clr4 mRNA (log2, nomr. to act1)
1
0
-1
-2
-3
WT
mst2Δepe1Δ
D
epe1Δ mst2deg
thiamine + NAA
-
+
IB: FLAG
*
Total protein
E
epe1Δ
mst2deg
+Mst2
-Mst2
1cm
F
epe1Δ
mst2deg
colony size (pixels2)
800
600
400
200
0
+Mst2
-Mst2
G
qRT-PCR
(-Epe1, 5 days)
clr4+
clr4 mRNA relative to N.S. (log2, norm. to tub1)
0
-1
-2
-3
mst2deg epe1+
mst2deg epe1Δ
mst2Δ epe1deg

Supp. 1

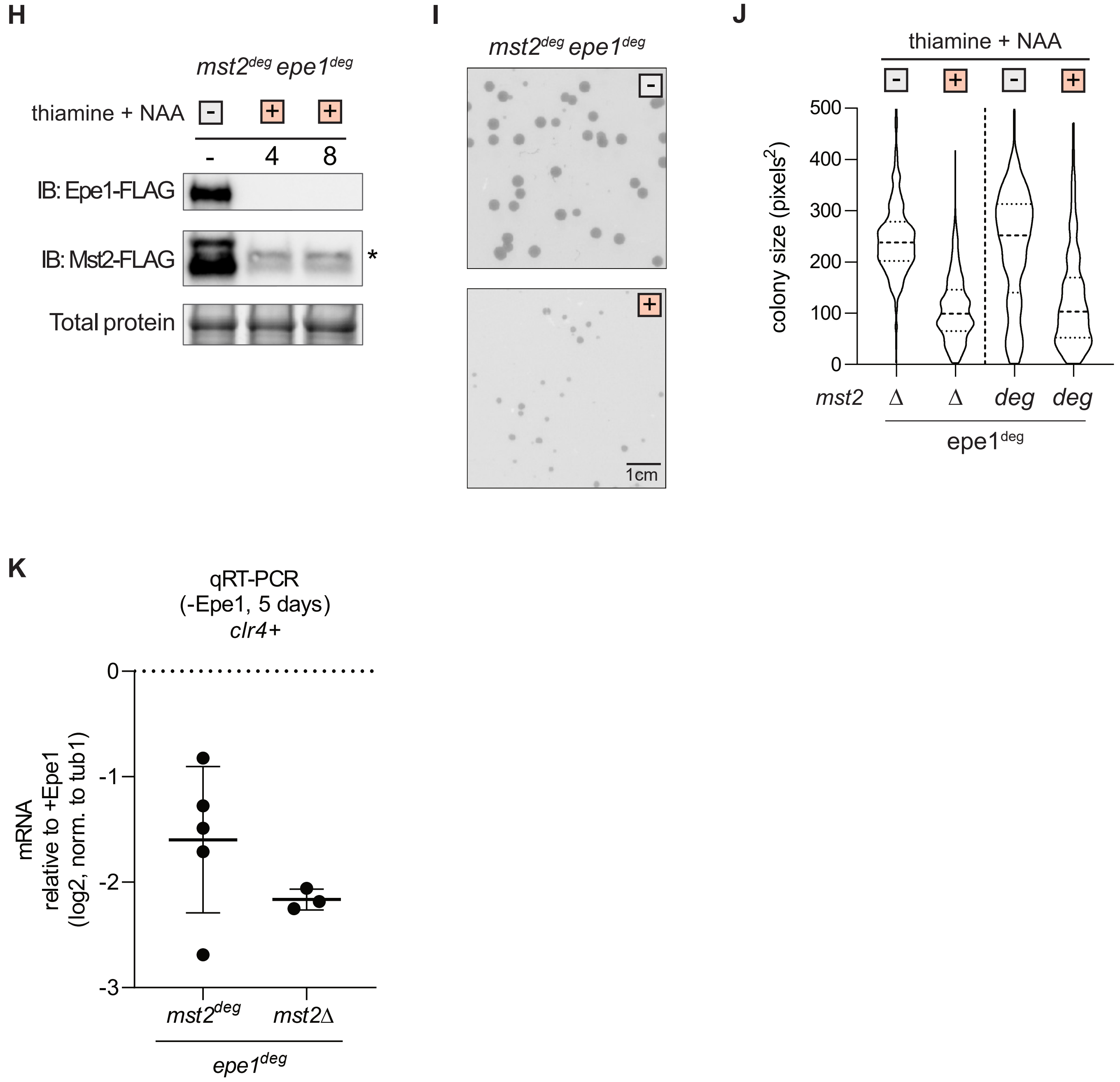

H
mst2deg epe1deg
thiamine + NAA
-
+
+
-
4
8
IB: Epe1-FLAG
IB: Mst2-FLAG
*
Total protein
I
mst2deg epe1deg
-
+
1cm
J
thiamine + NAA
-
+
-
+
500
400
300
200
100
0
colony size (pixels2)
mst2
Δ
Δ
deg
deg
epe1deg
K
qRT-PCR
(-Epe1, 5 days)
clr4+
0
-1
-2
-3
mRNA
relative to +Epe1
(log2, norm. to tub1)
mst2deg
mst2Δ
epe1deg

**Supplemental Figure 1**

(A) Example of a tetrad dissection of a meiotic cross between *mst2*Δ and *epe1*Δ cells. Picture was taken after 6 days of growth from tetrad containing all four genotype permutations.

(B) H3K9me2 ChIP-qPCR measuring enrichment at the *clr4+* locus. *mst2*Δ *epe1*Δ cells were all produced by meiotic crosses and grown for one week.

(C) *clr4*+ mRNA expression measured by qRT-PCR.

(D) Western blot for Mst2-3xFLAG-AID in *epe1*Δ $mst2^{deg}$. Media type is indicated with either a white box for no treatment, or an orange box for media with 15µM thiamine and 500µM NAA. Total protein levels are shown in the lower panel. Asterisk indicates nonspecific FLAG band, calibrated from samples without Mst2-3xFLAG-AID.

(E) Examples of *epe1*Δ $mst2^{deg}$ *S.pombe* colonies on solid media after three days of growth. Media type is indicated with either a white box for no treatment, or an orange box for media with 15µM thiamine and 500µM NAA. Image colors are inverted to highlight cell colonies.

(F) Colony size distribution, in pixel area, for *epe1*Δ $mst2^{deg}$ under different genetic backgrounds and growth conditions. Cell size quantified after five days of growth. Mean and st. dev of distributions in pixels$^2$: *epe1*Δ $mst2^{deg}$ no treatment (258.6 ± 87.5), thiamine and NAA (178.9 ± 71.6)

(G) *clr4*+ mRNA expression measured by qRT-PCR after five days of treatment with 15µM thiamine and 500µM NAA. Log2 fold-change expression of mRNA is relative to mRNA expression without thiamine and NAA. Error bars represent standard deviation, N=3.

(H) Western blot for Mst2-3xFLAG-AID and Epe1-3xFLAG-AID in $epe1^{deg}mst2^{deg}$. Media type is indicated with either a white box for no treatment, or an orange box for media with 15µM thiamine and 500µM NAA. Total protein levels are shown in the lower panel. Asterisk indicates nonspecific FLAG band, calibrated from samples without Mst2-3xFLAG-AID.

(I) Examples of $mst2^{deg}$ $epe1^{deg}$ *S.pombe* colonies on solid media after three days of growth. Media type is indicated with either a white box for no treatment, or an orange box for media with 15µM thiamine and 500µM NAA. Image colors are inverted to highlight cell colonies.

(J) Colony size distribution, in pixel area, for $mst2^{deg}$ $epe1^{deg}$ under different growth conditions. Cell size quantified after five days of growth. Distributions of *mst2*Δ $epe1^{deg}$ colony sizes are shown for comparison. Mean and st. dev of distributions in pixels$^2$: $mst2^{deg}$ $epe1^{deg}$ no treatment (230.3 ± 112.9), thiamine and NAA (127.0 ± 97.6)

(K) *clr4*+ mRNA expression measured by qRT-PCR after five days of treatment with 15µM thiamine and 500µM NAA. Log2 fold-change expression of mRNA is relative to mRNA expression without thiamine and NAA. Error bars represent standard deviation, N=3,5.

Supp. 2

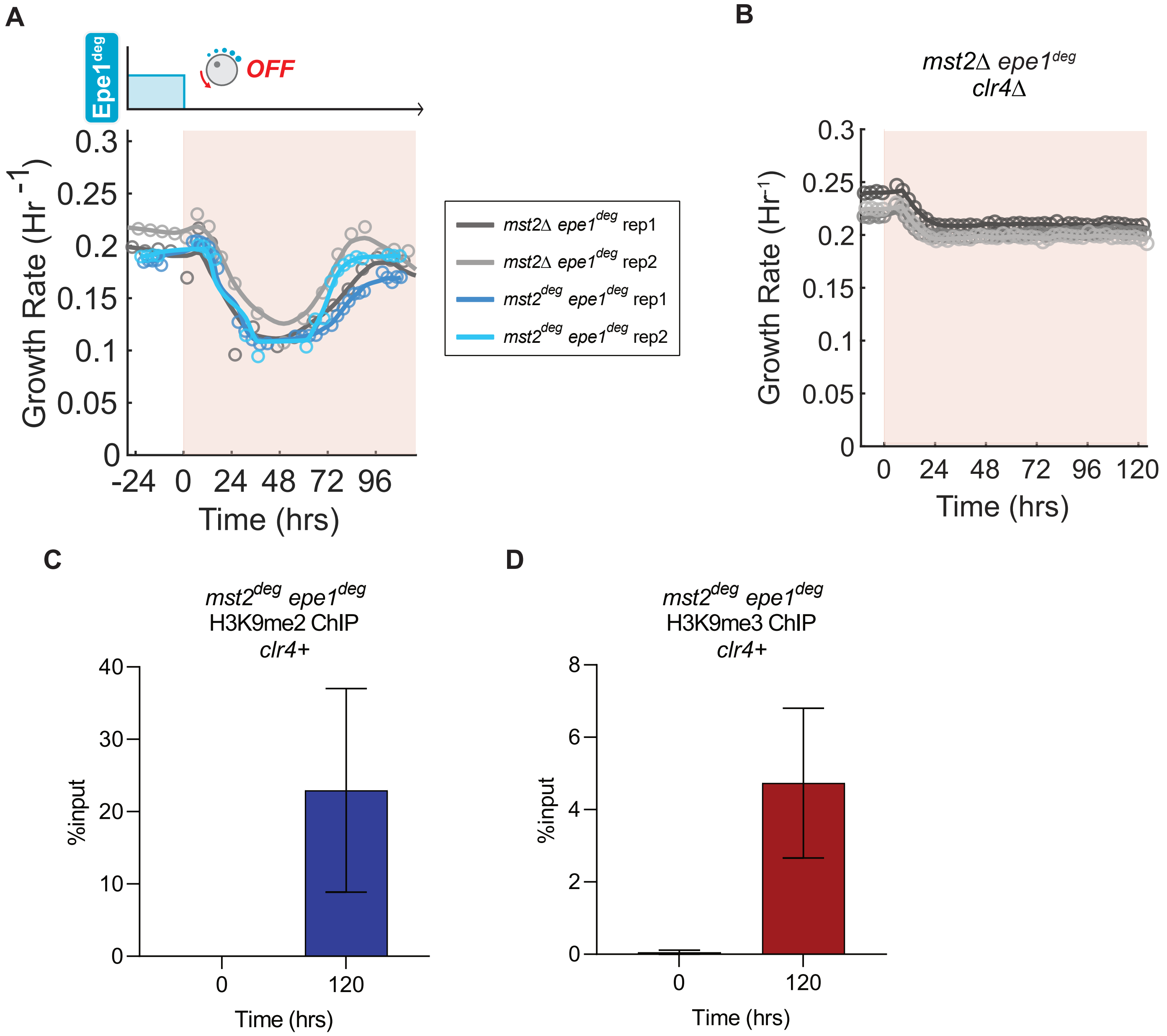
A
Epe1deg
OFF
Growth Rate (Hr -1)
0.3
0.25
0.2
0.15
0.1
0.05
0
-24 0 24 48 72 96
Time (hrs)
mst2Δ epe1deg rep1
mst2Δ epe1deg rep2
mst2deg epe1deg rep1
mst2deg epe1deg rep2
B
mst2Δ epe1deg
clr4Δ
Growth Rate (Hr-1)
0.3
0.25
0.2
0.15
0.1
0.05
0
0 24 48 72 96 120
Time (hrs)
C
mst2deg epe1deg
H3K9me2 ChIP
clr4+
%input
40
30
20
10
0
0
120
Time (hrs)
D
mst2deg epe1deg
H3K9me3 ChIP
clr4+
%input
8
6
4
2
0
0
120
Time (hrs)

E

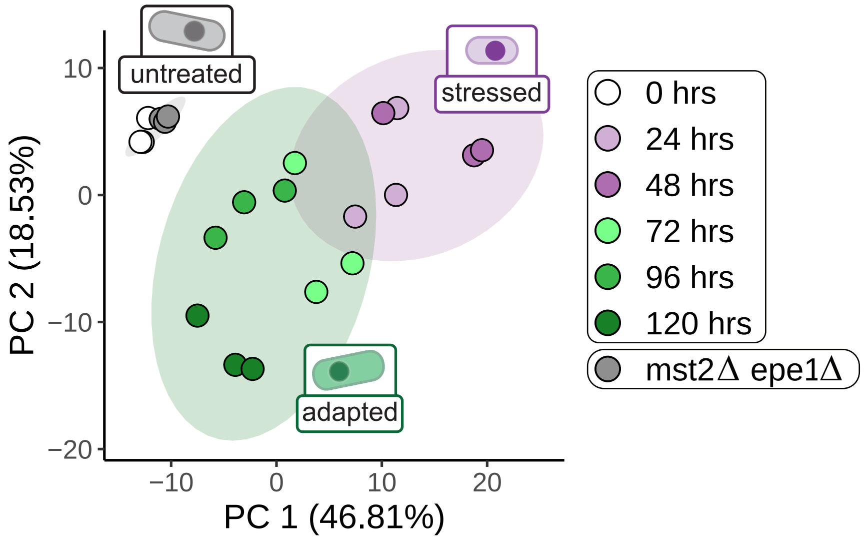
untreated
stressed
adapted
PC 2 (18.53%)
PC 1 (46.81%)
10
0
−10
−20
−10
0
10
20
0 hrs
24 hrs
48 hrs
72 hrs
96 hrs
120 hrs
mst2Δ epe1Δ

F

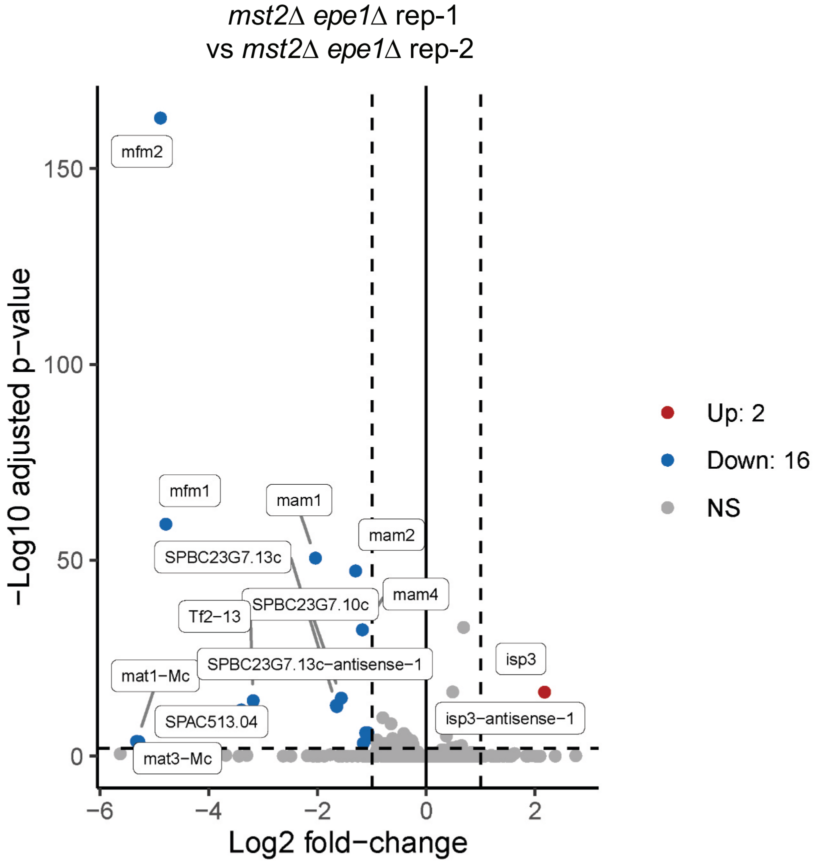
*mst2Δ epe1Δ* rep-1
vs *mst2Δ epe1Δ* rep-2
−Log10 adjusted p−value
Log2 fold−change
150
100
50
0
−6
−4
−2
0
2
mfm2
mfm1
mam1
mam2
SPBC23G7.13c
SPBC23G7.10c
mam4
Tf2−13
SPBC23G7.13c−antisense−1
isp3
mat1−Mc
SPAC513.04
isp3−antisense−1
mat3−Mc
Up: 2
Down: 16
NS

G

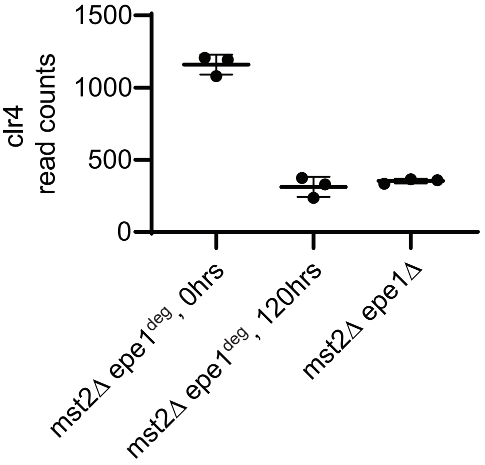
clr4
read counts
1500
1000
500
0
mst2Δ epe1deg, 0hrs
mst2Δ epe1deg, 120hrs
mst2Δ epe1Δ

H

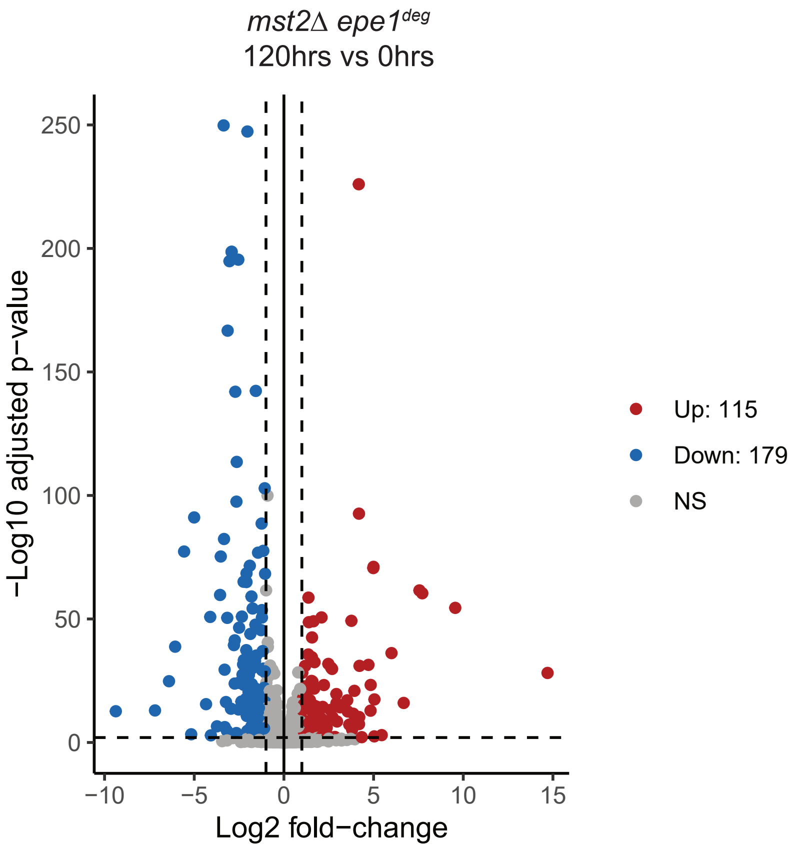


J

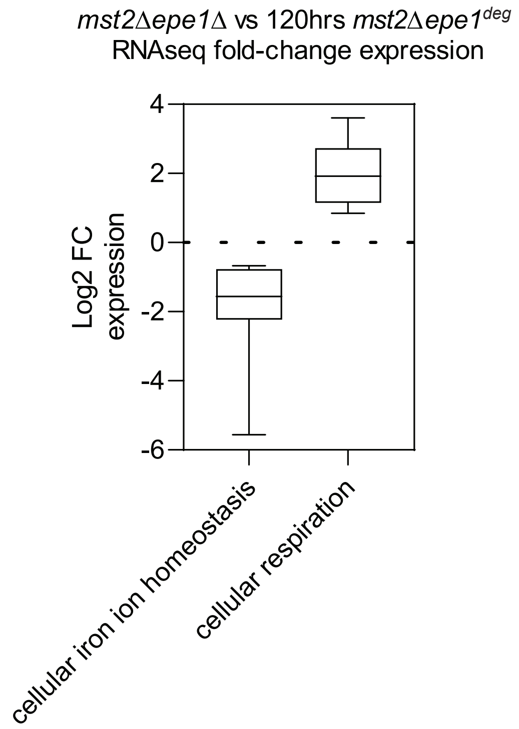


I

Upregulated genes at 120hrs vs 0hrs (Selected GO terms)

| Feature | Name | Frequency at 120hrs | Frequency in genome | Corrected Pvalue (-log10) |
|---|---|---|---|---|
| Biological Process | cellular iron ion homeostasis<br>response to iron ion starvation<br>cation transport | 6.95%<br>5.21%<br>10.43% | 0.27%<br>0.11%<br>1.87% | 6.54<br>6.33<br>3.61 |
| Gene Expression | caffeine and rapamycin induced<br>CESR induced | 18.26%<br>20.87% | 4.34%<br>7.72% | 5.11<br>3.12 |
| Molecular Functions | transmembrane transporter activity | 14.78% | 4.31% | 3.00 |

Downregulated genes at 120hrs vs 0hrs (Selected GO terms)

| Feature | Name | Frequency at 120hrs | Frequency in genome | Corrected Pvalue (-log10) |
|---|---|---|---|---|
| Biological Process | cellular respiration<br>ATP synthesis coupled electron transport | 8.38%<br>6.70% | 0.83%<br>0.39% | 8.32<br>9.31 |
| Cellular Component | mitochondrial respiratory chain | 6.15% | 0.40% | 7.80 |
| Transcript Features | ncRNA | 44.13% | 21.88% | 7.95 |

**Supplemental Figure 2**

(A) Real-time monitoring of growth rates of *mst2* $^{deg}$ *epe1* $^{deg}$ in eVOLVER, both replicates shown in blue. Treatment with 15µM thiamine and 500µM NAA was initiated at t=0hrs. Individual trendlines indicate replicates (N=2). Orange shaded portion represents the time period during which Epe1 has been depleted. Growth of *mst2*∆ *epe1* $^{deg}$ shown in grey tracks for comparison.

(B) Real-time monitoring of growth rates of *mst2*∆ *epe1* $^{deg}$ *clr4*∆ in eVOLVER. Treatment with 15µM thiamine and 500µM NAA was initiated at t=0hrs. Individual trendlines indicate replicates (N=2). Orange shaded portion represents the time period during which Epe1 has been depleted.

(C) H3K9me2 ChIP-qPCR measured at the *clr4*+ locus as a function of time following treatment of *mst2* $^{deg}$ *epe1* $^{deg}$ cells with 15µM thiamine and 500µM NAA. The orange shaded portion represents the time period during which Epe1 has been depleted. Error bars represent standard deviation, N=2.

(D) H3K9me3 ChIP-qPCR measured at the *clr4*+ locus as a function of time following treatment of *mst2* $^{deg}$ *epe1* $^{deg}$ cells with 15µM thiamine and 500µM NAA. Orange shaded portion represents the time period during which Epe1 has been depleted. Error bars represent standard deviation, N=2.

(E) Time course PCA analysis of the regularized log transform of RNAseq normalized counts denoting different time points after treatment with15µM thiamine and 500µM NAA. Colors denote untreated, stress and adapted cell phases. N=3, ellipse level=0.9.

(F) RNAseq volcano plot comparing transcriptomes of two independent *mst2*∆ *epe1*∆ cultures. Samples were produced from distinct crosses. Significance cutoffs are |FC| ≥ 1 and AdjPvalue ≤ 0.01, N=3.

(G) Read counts for *clr4*+ mRNA in RNAseq libraries.

(H) RNAseq volcano plot comparing transcriptomes of *mst2*∆ *epe1* $^{deg}$ treated with thiamine and NAA for 120 hours, versus untreated *mst2*∆ *epe1* $^{deg}$. Significance cutoffs are |FC| ≥ 1 and AdjPvalue ≤ 0.01, N=3.

(I) Table displaying selected GO terms for genes up- and downregulated in adapted *mst2*∆ *epe1* $^{deg}$ cells, compared to untreated cells. GO significance cutoff was set as FDR p-value ≤ 0.01.

(J) Boxplots showing average fold change expression for genes in selected GO terms for meiotically produced *mst2*∆ *epe1*∆ versus *mst2*∆ *epe1* $^{deg}$ treated with thiamine and NAA for 120 hours.

Supp. 3

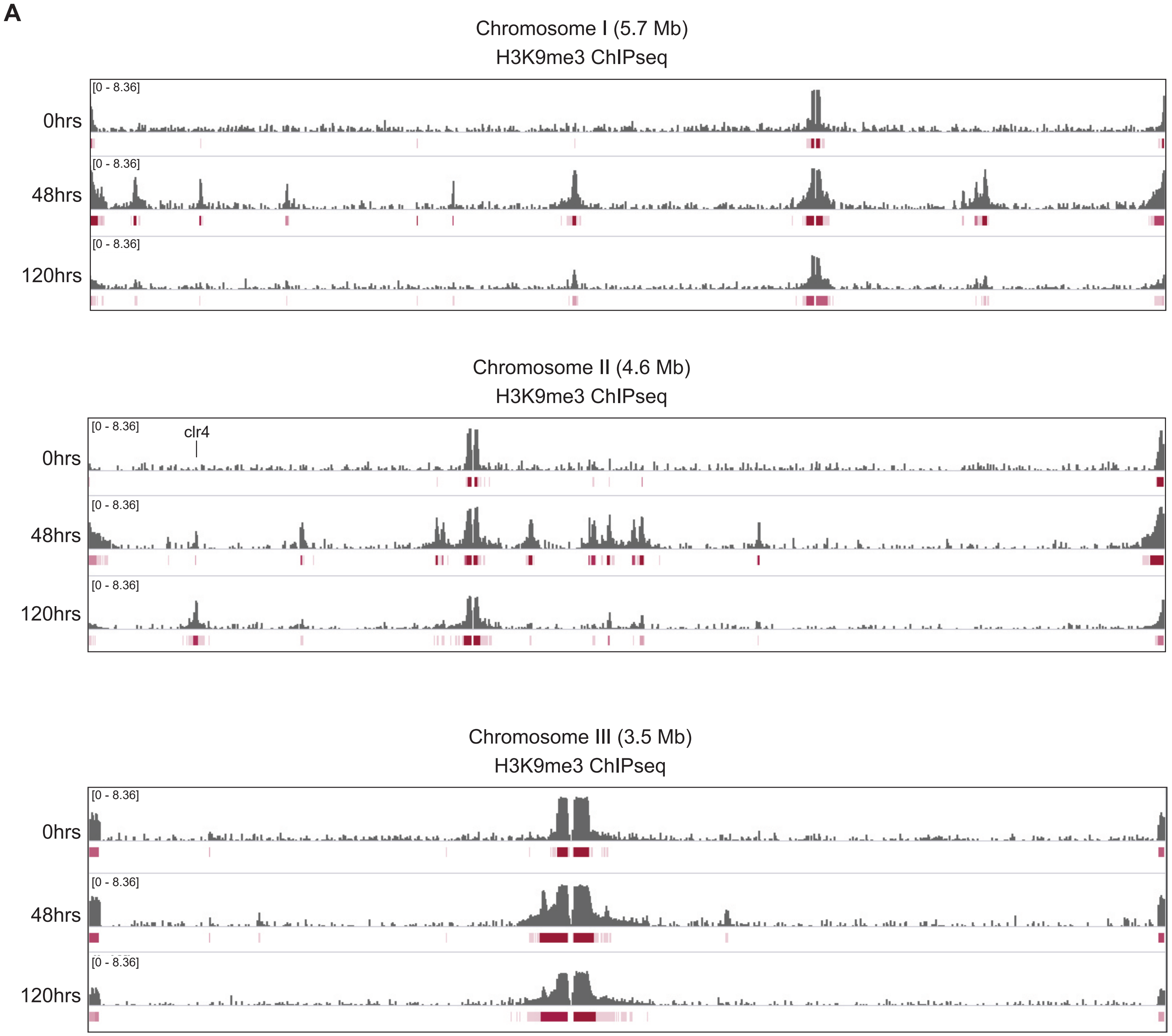

A
Chromosome I (5.7 Mb)
H3K9me3 ChIPseq
0hrs
48hrs
120hrs
[0 - 8.36]
Chromosome II (4.6 Mb)
H3K9me3 ChIPseq
clr4
0hrs
48hrs
120hrs
[0 - 8.36]
Chromosome III (3.5 Mb)
H3K9me3 ChIPseq
0hrs
48hrs
120hrs
[0 - 8.36]

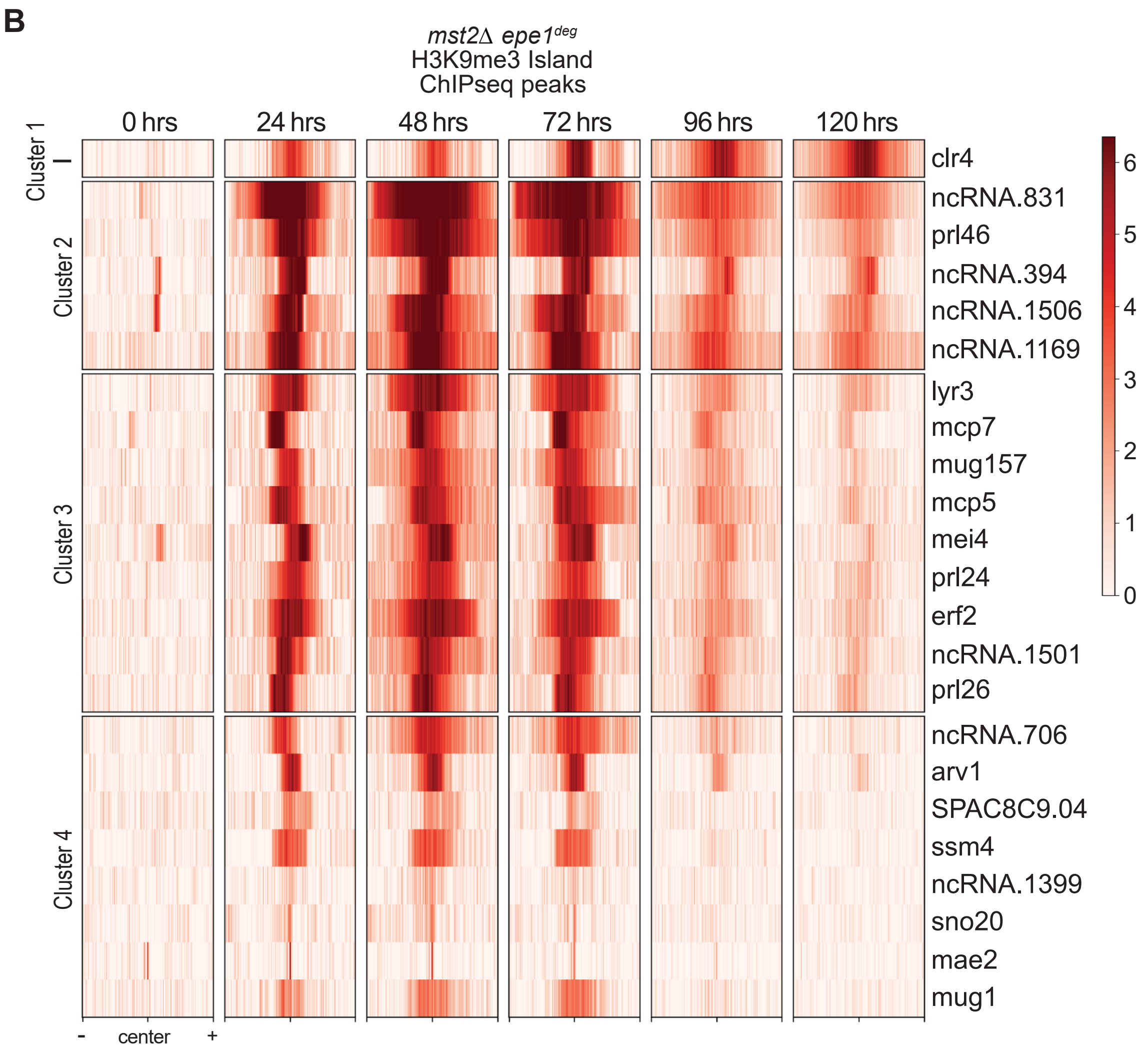
B
*mst2Δ epe1*[deg]
H3K9me3 Island
ChIPseq peaks
0 hrs
24 hrs
48 hrs
72 hrs
96 hrs
120 hrs
Cluster 1
Cluster 2
Cluster 3
Cluster 4
clr4
ncRNA.831
prl46
ncRNA.394
ncRNA.1506
ncRNA.1169
lyr3
mcp7
mug157
mcp5
mei4
prl24
erf2
ncRNA.1501
prl26
ncRNA.706
arv1
SPAC8C9.04
ssm4
ncRNA.1399
sno20
mae2
mug1
-
center
+
6
5
4
3
2
1
0

**C**

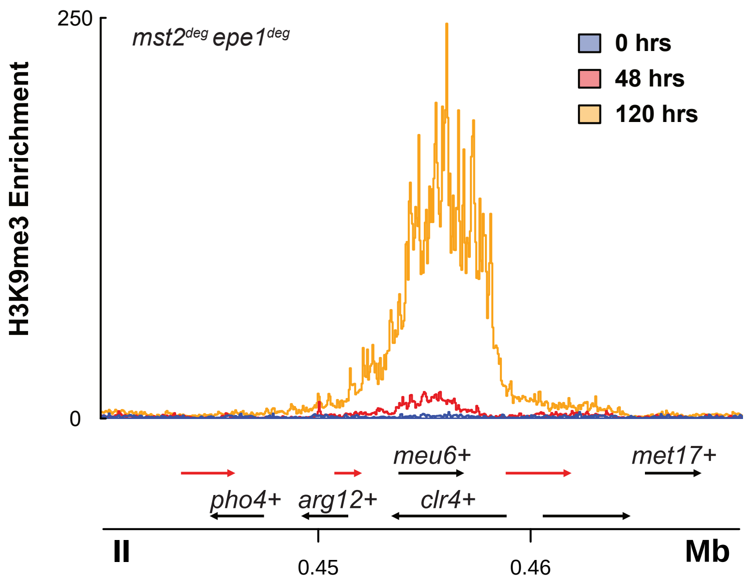
250
0
H3K9me3 Enrichment
mst2deg epe1deg
0 hrs
48 hrs
120 hrs
meu6+
met17+
pho4+
arg12+
clr4+
II
0.45
0.46
Mb

**D**

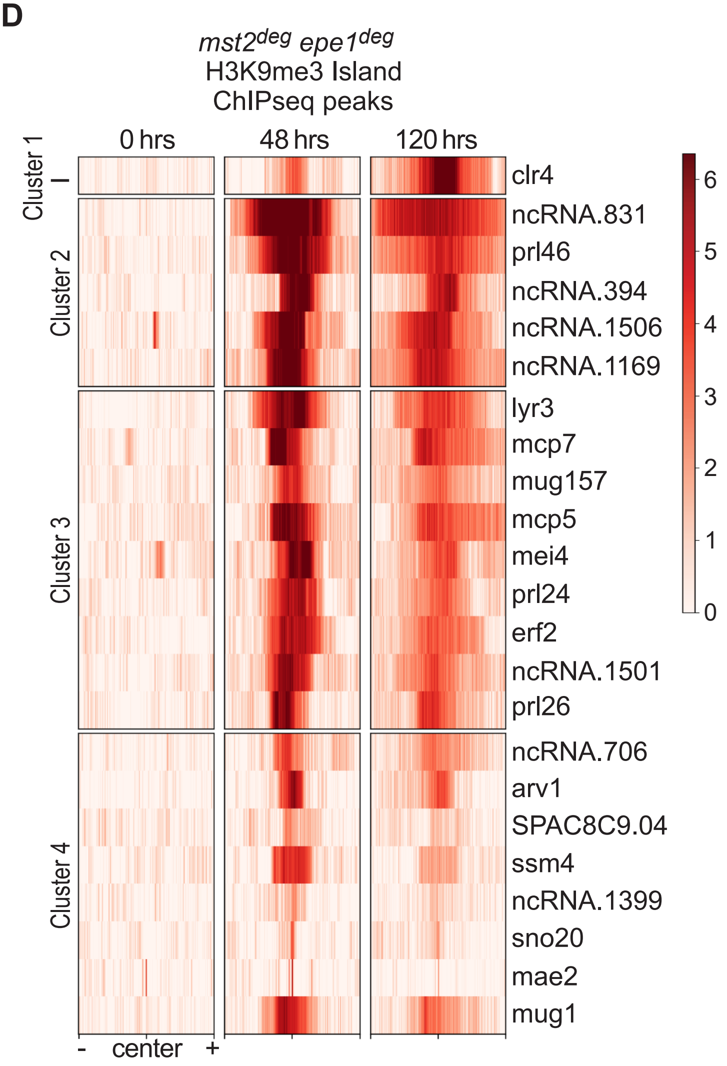
mst2deg epe1deg
H3K9me3 Island
ChIPseq peaks
0 hrs
48 hrs
120 hrs
Cluster 1
Cluster 2
Cluster 3
Cluster 4
clr4
ncRNA.831
prl46
ncRNA.394
ncRNA.1506
ncRNA.1169
lyr3
mcp7
mug157
mcp5
mei4
prl24
erf2
ncRNA.1501
prl26
ncRNA.706
arv1
SPAC8C9.04
ssm4
ncRNA.1399
sno20
mae2
mug1
- center +
6
5
4
3
2
1
0

Supp. 3

E

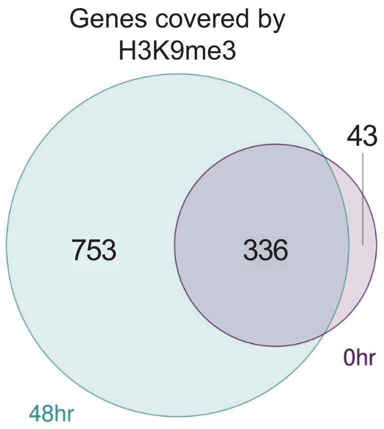
Genes covered by H3K9me3
43
753
336
0hr
48hr

F

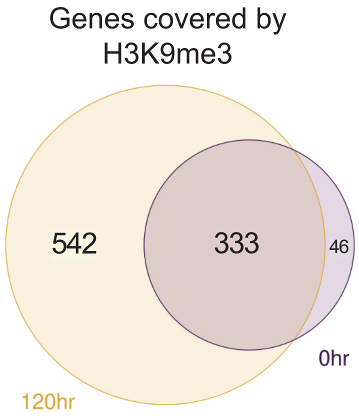
Genes covered by H3K9me3
542
333
46
0hr
120hr

G

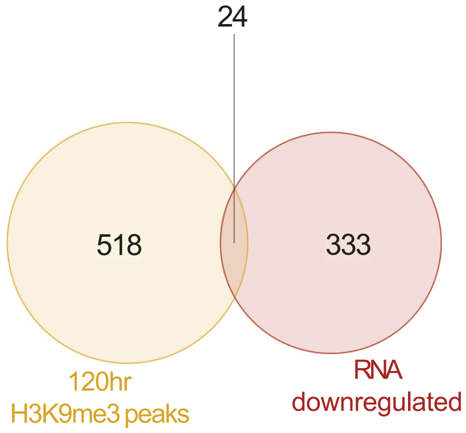
24
518
333
120hr H3K9me3 peaks
RNA downregulated

H

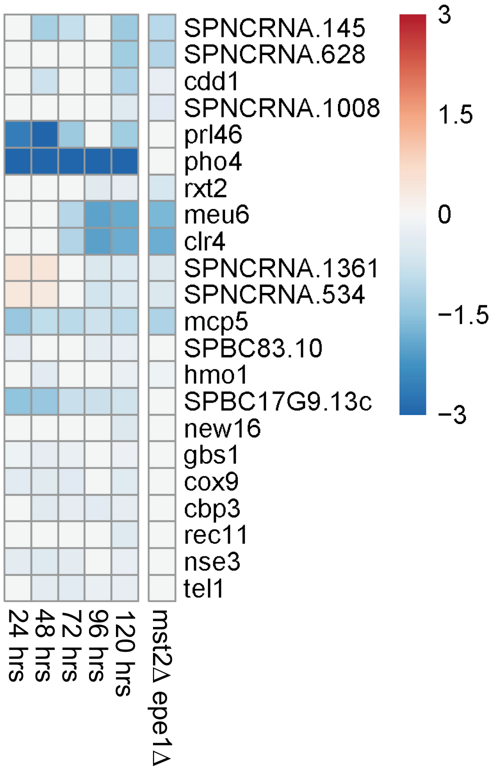
SPNCRNA.145
SPNCRNA.628
cdd1
SPNCRNA.1008
prl46
pho4
rxt2
meu6
clr4
SPNCRNA.1361
SPNCRNA.534
mcp5
SPBC83.10
hmo1
SPBC17G9.13c
new16
gbs1
cox9
cbp3
rec11
nse3
tel1
24 hrs
48 hrs
72 hrs
96 hrs
120 hrs
mst2Δ epe1Δ
3
1.5
0
−1.5
−3

**Supplemental Figure 3**

(A) H3K9me3 ChIP-seq tracks of the S. pombe genome. Enrichment is shown in log2 fold change of IP normalized to input. Time of Epe1 depletion is indicated on the left side of each track. Peaks identified are denoted in red below each track. The clr4+ gene locus is specifically highlighted. One of two ChIP-seq replicates is shown in this figure.

(B) H3K9me3 islands during heterochromatin misregulation at 0-120hrs of Epe1 depletion in *mst2*Δ *epe1*$^{deg}$. Peaks are centered in a 24kb window and are clustered by K-means clustering from figure 3E.

(C) H3K9me3 ChIP-seq enrichment centered on *clr4*+ for the indicated time points in *mst2*$^{deg}$ *epe1*$^{deg}$. Genomic tracks below show coding transcripts in black, non-coding transcripts in red.

(D) Heatmap showing loci of clustered H3K9me3 islands, originally identified in *mst2*Δ *epe1*$^{deg}$, in *mst2*$^{deg}$ *epe1*$^{deg}$. Peaks are centered in a 24kb window and are clustered by K-means clustering from figure 3E.

(E) Venn diagram depicting genes covered by H3K9me3 peaks at 0 hours and 48 hours of Epe1 depletion.

(F) Venn diagram depicting genes covered by H3K9me3 peaks at 0 hours and 120 hours of Epe1 depletion.

(G) Venn diagram depicting genes that are downregulated (AdjPval ≤ 0.01) by 120 hours after Epe1 depletion, overlapped with genes marked by H3K9me3 selectively at 120 hours.

(H) Heatmap depicting differential expression of the 24 genes covered by H3K9me3 peaks and are downregulated after 120 hours after Epe1 depletion. Changes in expression are log2 fold change relative to relative to untreated *mst2*Δ *epe1*$^{deg}$ cells. (AdjPval ≤ 0.01)

Supp. 4

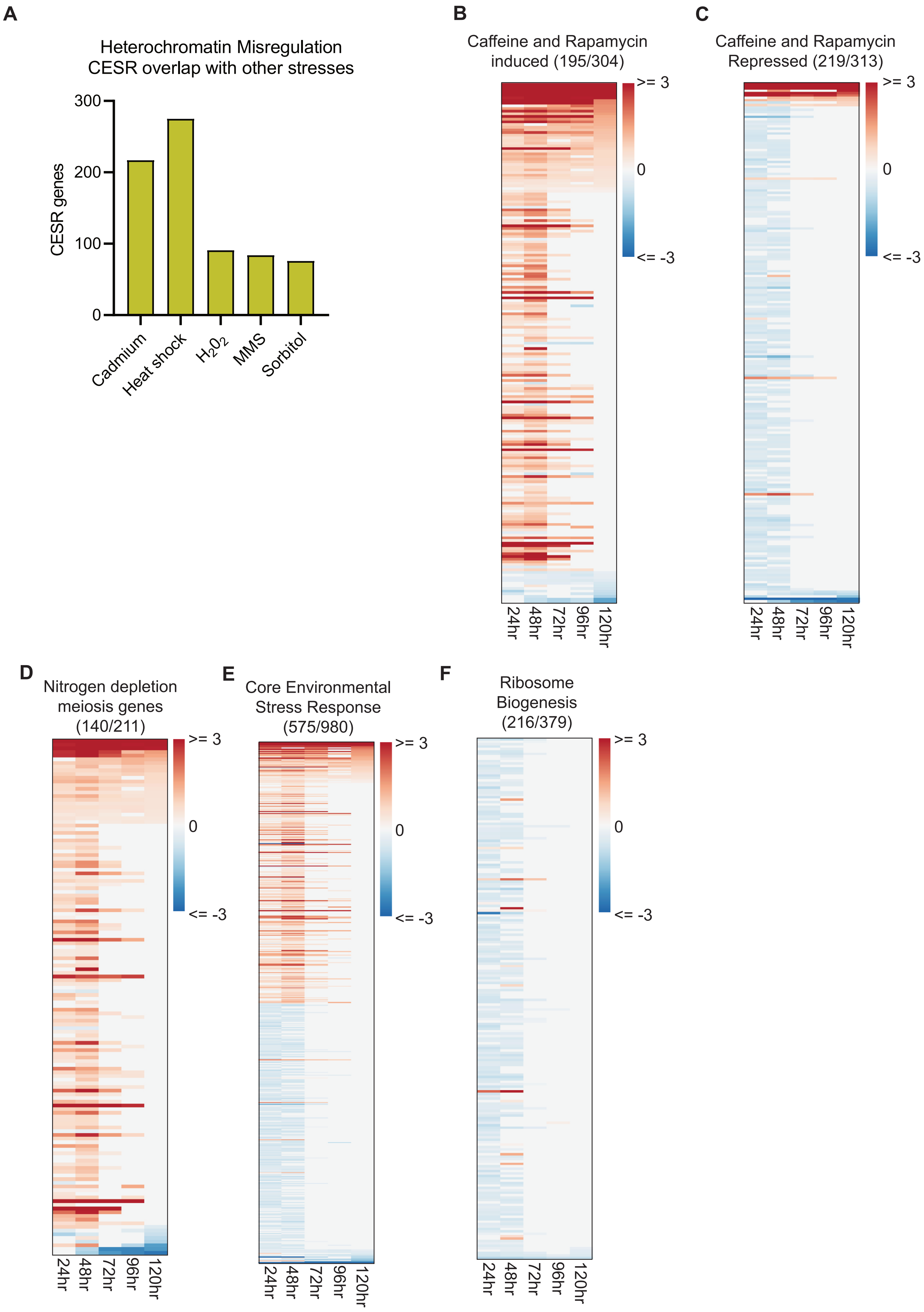
A
Heterochromatin Misregulation
CESR overlap with other stresses
CESR genes
300
200
100
0
Cadmium
Heat shock
H2O2
MMS
Sorbitol
B
Caffeine and Rapamycin
induced (195/304)
C
Caffeine and Rapamycin
Repressed (219/313)
D
Nitrogen depletion
meiosis genes
(140/211)
E
Core Environmental
Stress Response
(575/980)
F
Ribosome
Biogenesis
(216/379)
>= 3
0
<= -3
24hr
48hr
72hr
96hr
120hr

**Supplemental Figure 4**

(A) Bar graph indicating the overlap of CESR genes differentially expressed during stress phase of *mst2*Δ *epe1*$^{deg}$ that are similarly differentially expressed during indicated stresses.
(B) Heatmap of significant differential gene expression within the “Caffeine and Rapamycin induced” GO term relative to untreated *mst2*Δ *epe1*$^{deg}$ cells. Heatmap consists of genes that are differentially expressed at least during one time point. Total number of genes is indicated above the heatmap, significance cutoff of AdjPval ≤ 0.01.
(C) Heatmap of significant differential gene expression within the “Caffeine and Rapamycin repressed” GO term relative to untreated *mst2*Δ *epe1*$^{deg}$ cells. Heatmap consists of genes that are differentially expressed at least during one time point. Total number of genes is indicated above the heatmap, significance cutoff of AdjPval ≤ 0.01.
(D) Heatmap of significant differential gene expression within the “nitrogen depletion meiosis genes” GO term relative to untreated *mst2*Δ *epe1*$^{deg}$ cells. Heatmap consists of genes that are differentially expressed at least during one time point. Total number of genes is indicated above the heatmap, significance cutoff of AdjPval ≤ 0.01.
(E) Heatmap of significant differential gene expression within the “Core Environmental Stress Response” GO term relative to untreated *mst2*Δ *epe1*$^{deg}$ cells. Heatmap consists of genes that are differentially expressed at least during one time point. Total number of genes is indicated above the heatmap, significance cutoff of AdjPval ≤ 0.01.
(F) Heatmap of significant differential gene expression within the “Ribosome Biogenesis” GO term relative to untreated *mst2*Δ *epe1*$^{deg}$ cells. Heatmap consists of genes that are differentially expressed at least during one time point. Total number of genes is indicated above the heatmap, significance cutoff of AdjPval ≤ 0.01.

Supp Figure 5

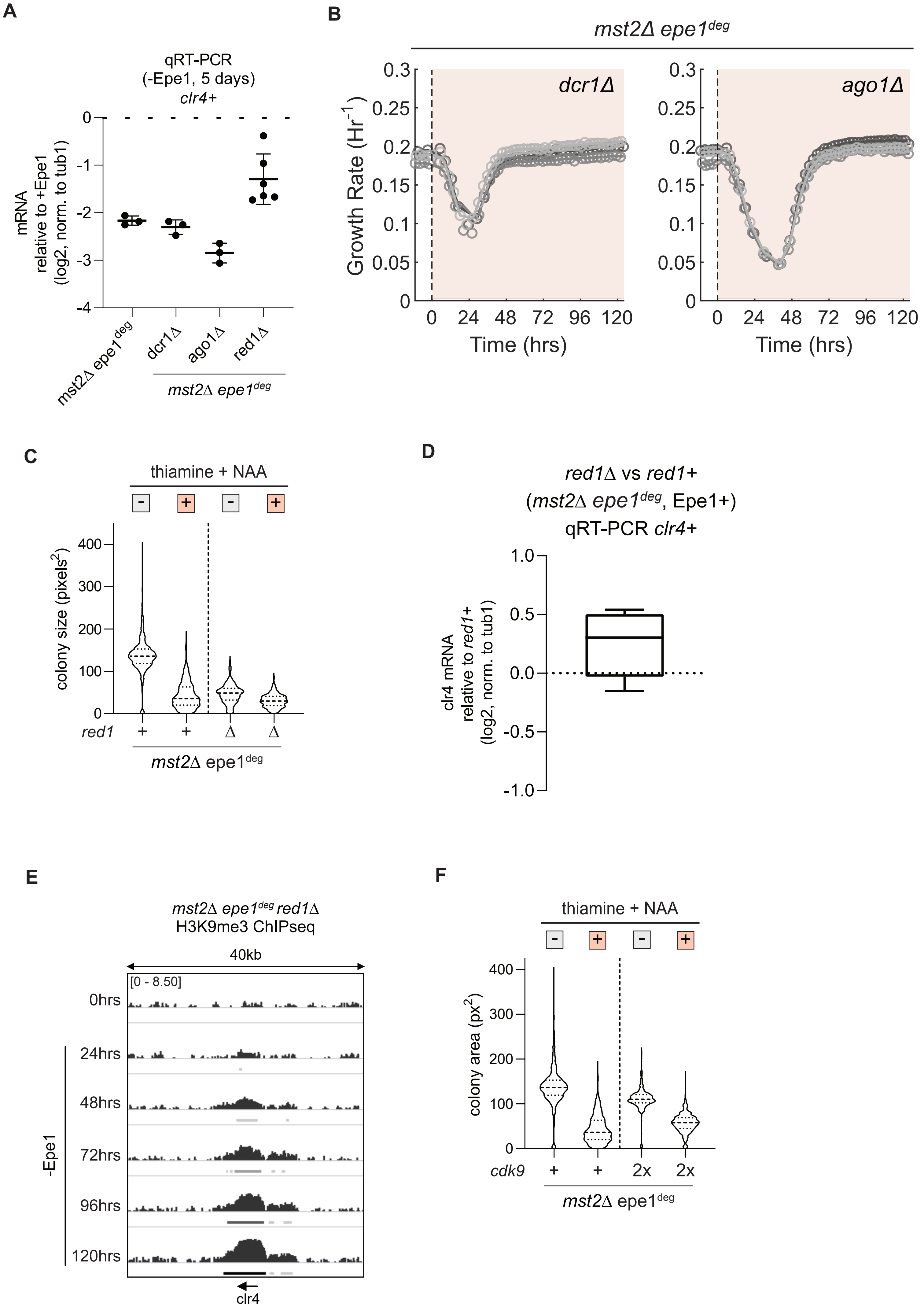

**Supplemental Figure 5**

(A) *clr4*+ mRNA expression measured by qRT-PCR after five days of treatment with 15µM thiamine and 500µM NAA. Log2 fold-change expression of mRNA is relative to mRNA expression without thiamine and NAA. Error bars represent standard deviation, N=3 or 6.

(B) Real-time monitoring of population growth rates of indicated genomic backgrounds cultured in eVOLVER. Treatment with 15µM thiamine and 500µM NAA was initiated at t=0hrs. Individual trendlines indicate replicates (N=2). Orange shaded portion represents the time period during which Epe1 has been depleted.

(C) Colony size distribution, in pixel area, for *mst2*Δ *epe1*$^{deg}$ *red1*Δ under different growth conditions. Cell size quantified after three days of growth. Distributions of *mst2*Δ *epe1*$^{deg}$ colony sizes are shown for comparison. Mean and st. dev of distributions in pixels$^2$: *mst2*Δ *epe1*$^{deg}$ *red1*Δ no treatment (47.3 ± 22.8), thiamine and NAA (30.7 ± 15.9)

(D) *clr4*+ mRNA expression measured by qRT-PCR in *mst2*Δ *epe1*$^{deg}$ and *mst2*Δ *epe1*$^{deg}$ *red1*Δ cells grown in EMMC. Log2 fold-change expression of mRNA is relative to mRNA expression without thiamine and NAA.

(E) H3K9me3 ChIP-seq (40kb window) centered at the *clr4*+ gene locus after five days of Epe1 depletion in *mst2*Δ *epe1*$^{deg}$ *red1*Δ cells. Enrichment is shown in log2 fold change of IP normalized to input. Epe1 was depleted within the window of 0-120 hours. The *clr4*+ gene locus is specifically highlighted.

(F) Colony size distribution, in pixel area, for *mst2*Δ *epe1*$^{deg}$ *leu1:cdk9*+ under different growth conditions. Cell size quantified after three days of growth. Distributions of *mst2*Δ *epe1*$^{deg}$ colony sizes are shown for comparison. Mean and st. dev of distributions in pixels$^2$: *mst2*Δ *epe1*$^{deg}$ *leu1:cdk9*+ no treatment (111.6 ± 25.3), thiamine and NAA (57.2 ± 21.8)

Supp 6

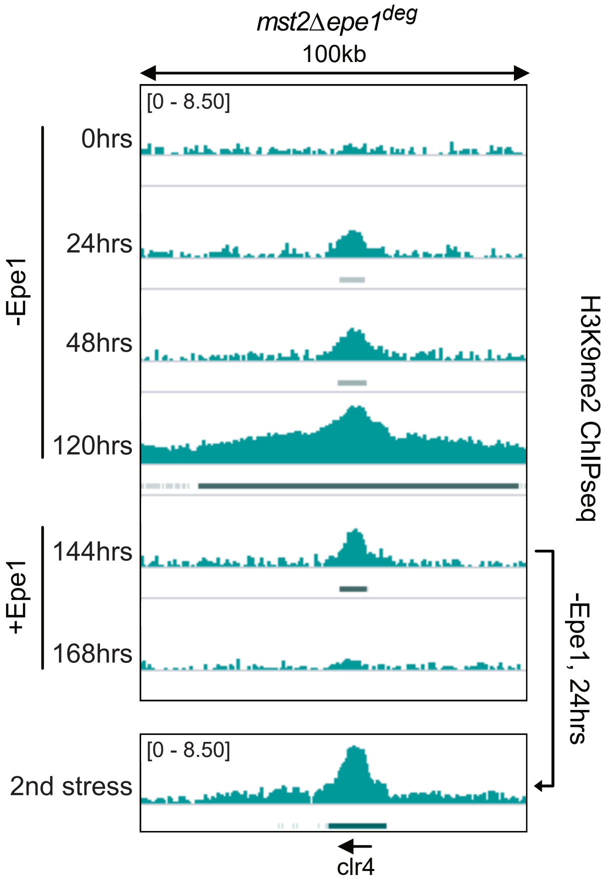
mst2Δepe1deg
100kb
[0 - 8.50]
0hrs
24hrs
48hrs
120hrs
144hrs
168hrs
-Epe1
+Epe1
H3K9me2 ChIPseq
-Epe1, 24hrs
[0 - 8.50]
2nd stress
clr4

**Supplemental Figure 6**

H3K9me2 ChIP-seq tracks in a 100kb window centered at the *clr4+* gene locus after five days of Epe1 depletion in an *mst2*Δ background. Enrichment is shown in log2 fold change of IP normalized to input. Epe1 was depleted within the window of 0-120 hours. After 120 hours, Epe1 was expressed for a recovery period of 24, 48, or 72 hours. Recovered populations were then put under a second stress by depleting Epe1 for another 24 hours.. Peaks identified are denoted in red below each track. The *clr4+* gene locus is specifically highlighted.

Supp. Figure 7

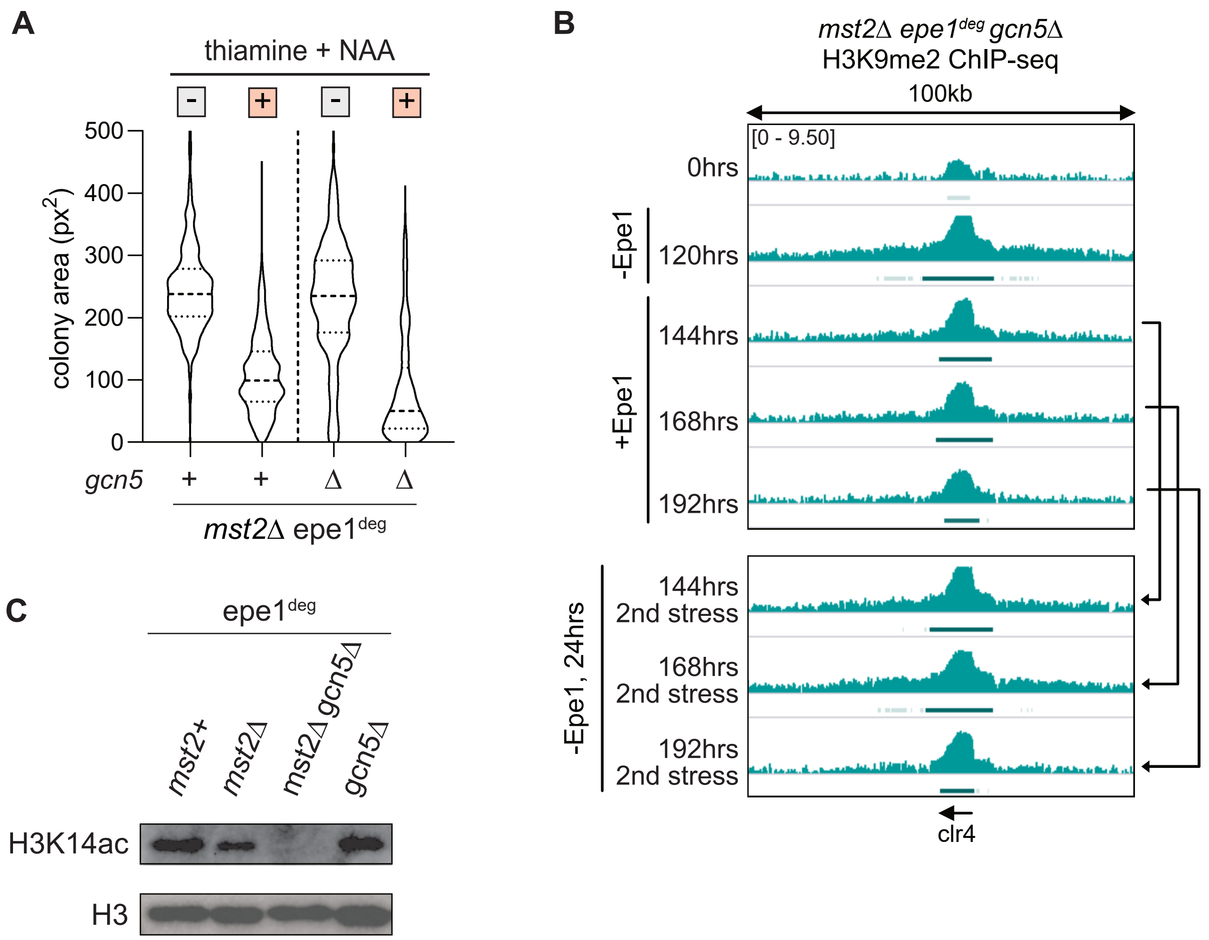

Supp. Figure 7

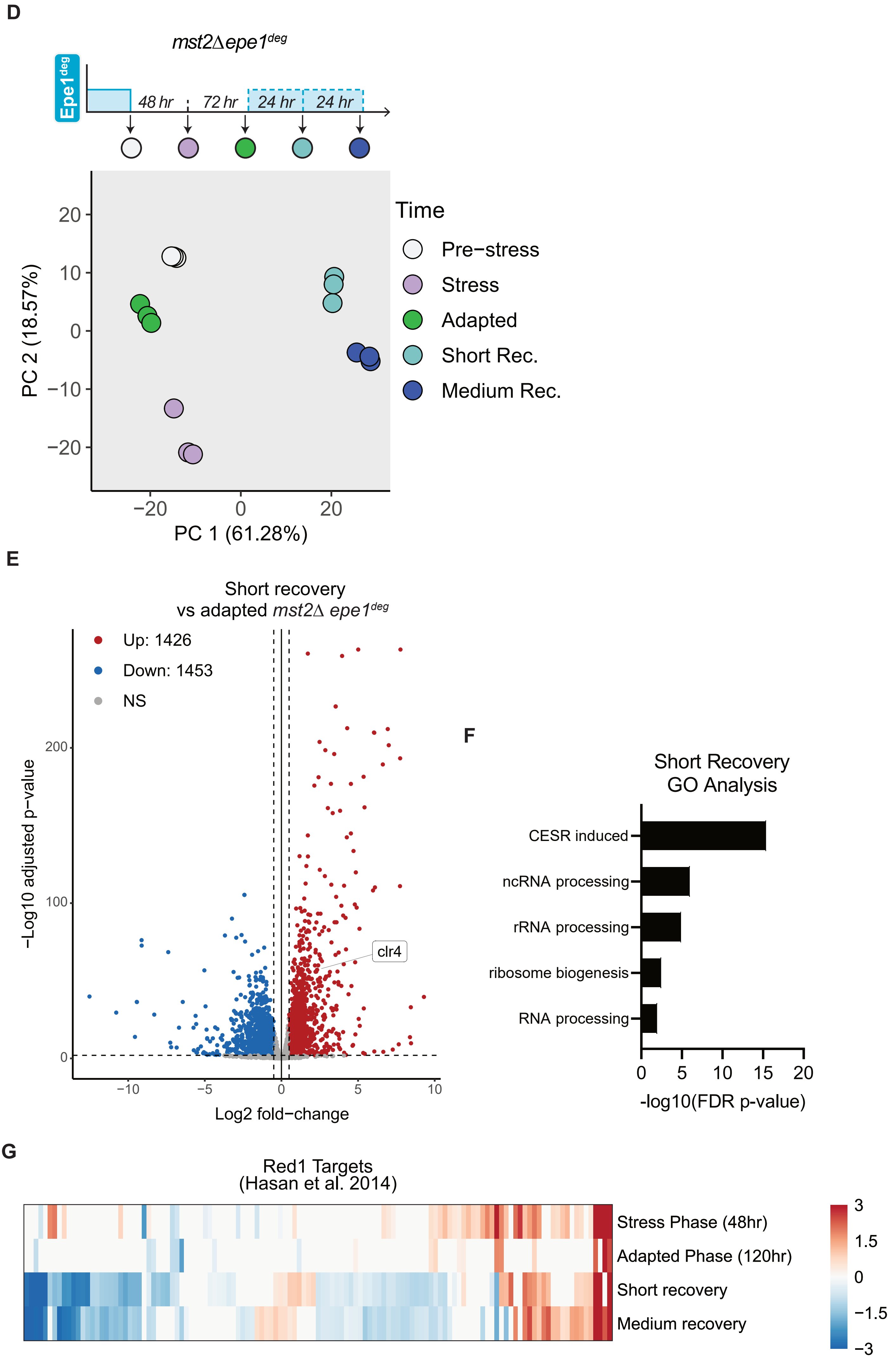

**Supplemental Figure 7**

(A) Colony size distribution, in pixel area, for *mst2*Δ *epe1$^{deg}$* *gcn5*Δ under different growth conditions. Cell size quantified after five days of growth. Distributions of *mst2*Δ *epe1$^{deg}$* colony sizes are shown for comparison. Mean and st. dev of distributions in pixels$^2$: *mst2*Δ *epe1$^{deg}$* *gcn5*Δ no treatment (202.8 ± 98.3), thiamine and NAA (72.2 ± 67.6)

(B) H3K9me2 ChIP-seq tracks in a 100kb window centered at the *clr4*+ gene locus after five days of Epe1 depletion in an *mst2*Δ *epe1$^{deg}$* *gcn5*Δ background. Enrichment is shown in log2 fold change of IP normalized to input. Epe1 was depleted within the window of 0-120 hours. After 120 hours, Epe1 was expressed for a recovery period of 24, 48, or 72 hours. Recovered populations were then put under a second stress by depleting Epe1 for another 24 hours. Peaks identified are denoted below each track. The *clr4*+ gene locus is specifically highlighted.

(C) Western blot comparing total H3K14ac levels in indicated genotypes. H3 is shown as a loading control.

(D) Time course PCA analysis of the regularized log transform of RNAseq normalized counts denoting different time points after treatment with15µM thiamine and 500µM NAA. Colors denote untreated, stress, adapted, and recovery cell phases. N=3, ellipse level=0.9. Epe1 expression and time points are shown above the figure.

(E) RNAseq volcano plot comparing transcriptomes of *mst2*Δ *epe1$^{deg}$* in short recovery versus the adapted phase. Significance cutoffs are |FC| ≥ 0.5 and AdjPvalue ≤ 0.01, N=3.

(F) Selected GO terms for transcripts differentially expressed (AdjPval ≤ 0.01) during short recovery, relative to untreated. GO significance cutoff was set as FDR p-value ≤ 0.01.

(G) Heatmap of Red1 target transcripts that are differentially expressed, relative to untreated *mst2*Δ *epe1$^{deg}$* cells. Heatmap consists of transcripts that are differentially expressed at least during one time point. 125 transcripts are shows, significance cutoff of AdjPval ≤ 0.01.

### Tables

#### T.1 *S.pombe* strains

| Strain no. | Genotype |
|---|---|
| SPYKR-477 | h- ade6::ade6+-adh15prom-skp1-OsTIR1-natMX6-adh15prom-skp1-AtTIR1-2NLS ura4-D18 |
| SPYKR-521 | h+ mst2::ura4+ ura4-D18 ade6-M210 leu1-32 |
| SPYKR-1133 | h- ade6::ade6+-adh15prom-skp1-OsTIR1-natMX6-adh15prom-skp1-AtTIR1-2NLS ura4-D18 mst2Δ::ura4 |
| SPYKR-1142 | h- ade6::ade6+-adh15prom-skp1-OsTIR1-natMX6-adh15prom-skp1-AtTIR1-2NLS ura4-D18 epe1Δ::bsd |
| SPYKR-1145 | h- leu1-32 ade6-M210 ura4-D18 his3-Dr epe1Δ::bsd |
| SPYKR-1183 | h- ade6::ade6+-adh15prom-skp1-OsTIR1-natMX6-adh15prom-skp1-AtTIR1-2NLS ura4-D18 nmt81-mst2-3xFLAG-AID-kanMx6 |
| SPYKR-1218 | h- ade6::ade6+-adh15prom-skp1-OsTIR1-natMX6-adh15prom-skp1-AtTIR1-2NLS ura4-D18 nmt81-mst2-3xFLAG-AID-kanMx6 epe1Δ::bsd |
| SPYKR-1265 | h- ade6::ade6+-adh15prom-skp1-OsTIR1-natMX6-adh15prom-skp1-AtTIR1-2NLS ura4-D18 nmt81-epe1-3xFLAG-AID-kanMX6 |
| SPYKR-1351 | h- ade6::ade6+-adh15prom-skp1-OsTIR1-natMX6-adh15prom-skp1-AtTIR1-2NLS ura4-D18 nmt81-epe1-3xFLAG-AID-kanMX6 mst2Δ::ura4 #1 |
| SPYKR-1492 | h- ade6::ade6+-adh15prom-skp1-OsTIR1-natMX6-adh15prom-skp1-AtTIR1-2NLS ura4-D18 nmt81-epe1-3xFLAG-AID-bsdMX6 mst2Δ::ura4 |
| SPYKR-1650 | h+ ura4-D18 leu1-32 ade6::ade6+-adh15prom-skp1-OsTIR1-natMX6-adh15prom-skp1-AtTIR1-2NLS nmt81-epe1-3xFLAG-AID-bsdMX6 mst2Δ::ura4+ dcr1Δ::hphMX6 |
| SPYKR-1673 | h- ade6::ade6+-adh15prom-skp1-OsTIR1-natMX6-adh15prom-skp1-AtTIR1-2NLS ura4-D18 nmt81-epe1-3xFLAG-AID-bsdMX6 mst2Δ::ura4 ago1Δ::kanMX6 |
| SPYKR-1732 | h- ade6::ade6+-adh15prom-skp1-OsTIR1-natMX6-adh15prom-skp1-AtTIR1-2NLS ura4-D18 nmt81-epe1-3xFLAG-AID-bsdMX6 mst2Δ::ura4 sty1Δ::kanMX6 |
| SPYKR-2019 | h- ade6::ade6+-adh15prom-skp1-OsTIR1-natMX6-adh15prom-skp1-AtTIR1-2NLS ura4-D18 nmt81-epe1-3xFLAG-AID-bsdMX6 mst2Δ::ura4 red1Δ::hphMX6 |
| SPYKR-2109 | h- ade6::ade6+-adh15prom-skp1-OsTIR1-natMX6-adh15prom-skp1-AtTIR1-2NLS ura4-D18 nmt81-epe1-3xFLAG-AID-bsdMX6 mst2Δ::ura4 gcn5Δ::G418 |
| SPYKR-3032 | h- ade6::ade6+-adh15prom-skp1-OsTIR1-natMX6-adh15prom-skp1-AtTIR1-2NLS ura4-D18 nmt81-epe1-3xFLAG-AID-bsdMX6 nmt81-mst2-3xFLAG-AID-kanMX6 |
| SPYKR-3230 | h- ade6::ade6+-adh15prom-skp1-OsTIR1-natMX6-adh15prom-skp1-AtTIR1-2NLS ura4-D18 nmt81-epe1-3xFLAG-AID-bsdMX6 mst2Δ::mst2+ gcn5Δ::G418 |
| SPYKR-MC1 | h+ mst2::ura4+ leu1-32 ade6-M210 ura4-D18 his3? epe1Δ::bsd |
| SPYKR-MC2 | h- mst2::ura4+ leu1-32 ade6-M210 ura4-D18 his3? epe1Δ::bsd |

**T.2 Plasmids**

| Plasmid no. | Description |
|---|---|
| KRP-256 | pFA6a 3xflag AID IAA-17 degron kanMX6, Moazed lab |
| KRP-472 | pFA6a nmt81 3xflag AID IAA-17 degron kanMX6 |
| KRP-485 | pFA6a nmt81 mst2 3xflag AID IAA-17 degron |
| KRP-487 | pFA6a nmt81 epe1 3xflag AID IAA-17 degron |

### T.3 Primers

| Primer sequence | Notes |
|---|---|
| AACGCTTGGCCATGGAATACACG | tub1 qPCR forward primer |
| GAGAGGCGGTGATGGAAGAAACAAC | tub1 qPCR reverse primer |
| AACCCTCAGCTTTGGGTCTT | act1 qPCR forward primer |
| TTTGCATACGATCGGCAATA | act1 qPCR reverse primer |
| CAACATGAAGAAATGAAAGCGTATCG | epe1 qPCR forward primer |
| CTCTGGACCGAAGCCTGTGCC | epe1 qPCR reverse primer |
| ATGCGAGCAATCCACATCCA | clr4 qPCR forward primer |
| ATCATTCGTCGTCAGAGCGG | clr4 qPCR reverse primer |
| CGCTGCAGGTCGACGGATCCCCGGGAGATCTCGCCATAAAAGACAGAATAAG | amplify nmt81 promoter into pacI digested 256 plasmid - forward |
| ATCGTCGTCTTTATAATCAAAGATGTTAATTAAGATTTAACAAAGCGACTATAAGTCAGA | amplify nmt81 promoter into pacI digested 256 plasmid - reverse |
| CTTATAGTCGCTTTGTTAAATCTTAATTAAATGGATTCCTGGCTTGAATACG | FWD epe1 insertion into plasmid 471/472 pacI site |
| CTTATCATCGTCGTCTTTATAATCAAAGATACTAGCACCTCTGGACCGAAG | REV epe1 insertion into plasmid 471/472 pacI site |
| CTTATAGTCGCTTTGTTAAATCTTAATTAAATGCCGGCGACTATTTTGAA | FWD mst2 insertion into plasmid 471/472 pacI site |
| CTTATCATCGTCGTCTTTATAATCAAAGATAACGGAATCCAGATGATGAGAGTT | REV mst2 insertion into plasmid 471/472 pacI site |
| ATCTCCTTAGTTTGCATCGCAATTTTATAGTTACCTTTTTGCTAGTAAGCAATTAATTTTTG GGACTTTTAAGAGATCTCGCCATAAAAGACAGAATAAG | FWD epe1 amplify with nmt promoter |
| AAGGATTAAATGGGGCCATTAGGAAAACTATGAACGATCTGTAAATATAAACAATCTTTTT TTTTTATGTATAAGATCTCGCCATAAAAGACAGAATAAG | FWD mst2 amplify with nmt promoter |
| CGTCAAGACTGTCAAGGAGGGTATTC | REV insert nmt-gene cassette |

**T.4 Heterochromatin misregulation islands at 48hrs of Epe1 depletion**

| row | cluster | chrom | centered gene | 48hrs H3K9me3 peak start | 48hrs H3K9me3 peak end |
|---|---|---|---|---|---|
| 1 | 1 | II | clr4 | 451877 | 457826 |
| 2 | 2 | I | ncRNA.831 | 2510924 | 2533382 |
| 3 | 2 | I | prl46 | 4638558 | 4664225 |
| 4 | 2 | II | ncRNA.394 | 2192141 | 2202241 |
| 5 | 2 | II | ncRNA.1506 | 2329293 | 2346043 |
| 6 | 2 | III | ncRNA.1169 | 1029501 | 1047111 |
| 7 | 3 | I | lyr3 | 236009 | 252524 |
| 8 | 3 | I | mcp7 | 577626 | 587367 |
| 9 | 3 | I | mug157 | 4596991 | 4610271 |
| 10 | 3 | II | mcp5 | 895485 | 907436 |
| 11 | 3 | II | mei4 | 1466522 | 1477016 |
| 12 | 3 | II | prl24 | 1491884 | 1502976 |
| 13 | 3 | II | erf2 | 1860059 | 1876982 |
| 14 | 3 | II | ncRNA.1501 | 2297436 | 2309154 |
| 15 | 3 | II | prl26 | 2826386 | 2835892 |
| 16 | 4 | I | ncRNA.706 | 1023545 | 1035895 |
| 17 | 4 | I | arv1 | 1890359 | 1895878 |
| 18 | 4 | I | SPAC8C9.04 | 3648079 | 3649030 |
| 19 | 4 | I | ssm4 | 4533090 | 4540616 |
| 20 | 4 | II | ncRNA.1399 | 949525 | 949932 |
| 21 | 4 | II | sno20 | 2410271 | 2411017 |
| 22 | 4 | III | mae2 | 276537 | 276889 |
| 23 | 4 | III | mug1 | 1452148 | 1457902 |
